# Phase similarity between similar objects indicates representational merging across retrieval training but not sleep

**DOI:** 10.64898/2026.01.18.700222

**Authors:** Hayley Bree Caldwell, Alex Chatburn, Kurt Lushington, Simon Hanslmayr, Sebastian Michelmann

## Abstract

Retrieval training (i.e., cued recall) is theorised to induce rapid memory consolidation, similarly to sleep. Across consolidation, related neural representations become increasingly similar; yet, this representational change has never been directly compared between sleep and retrieval training. In this study, 30 subjects (27F, 18–34, *M*=22.17) completed four separate sessions in which they (1) learnt object-word pairs, followed by (2) immediate recognition testing, (3) one of four 120-min interventions (retrieval training, restudy, sleep, or wake), and (4) delayed recognition testing. We compared EEG phase similarity between similar and different objects to assess the time, frequency, and anatomical distribution of representational similarity across encoding (learning to immediate recognition), and each intervention (immediate to delayed recognition). We hypothesised that EEG phase patterns for similar objects would become more similar (i.e., representational merging) across retrieval training and sleep interventions, and predict a greater endorsement of similar-object lures. Indeed, we found increased representational similarity between similar objects across the encoding shift in the theta-band and occipital sources. Crucially, additional representational merging was only observed across the retrieval training intervention, in the alpha-band and parieto-occipital sources. Despite retrieval training leading to reduced performance in discriminating similar-objects lures, greater representational merging across retrieval training predicted greater discrimination of similar-object lures. Together, these findings suggest that sleep and retrieval training induce different memory transformations across the same timescale. Retrieval training may generally provoke rapid gist extraction, with greater neocortical integration supporting episodic discrimination. Conversely, sleep may selectively maintain task-relevant episodic and semantic details in the short-term.

## Introduction

One of the most important aspects of the brain’s memory system is its ability to efficiently restructure memory representations for their long-term access. This restructuring is achieved via memory consolidation and involves the semanticization and gradual integration of memories into neocortical networks, leaving them less dependent on the hippocampus [1,2]. Non-rapid eye movement (NREM) sleep, even as short as an afternoon nap, provides the best conditions for memory consolidation, resulting in gist-like representations of memories and a reduction of memory for contextual details [3–5]. However, recent evidence suggests that the same representational changes observed following sleep, can also be observed during wake following repeated retrieval training [6–9]. Whilst several studies have explored memory outcomes across sleep-based and retrieval-mediated memory consolidation, no study to our knowledge has compared the representational changes across both consolidation types. Additionally, existing methods that compare memory outcomes between sleep and retrieval training typically include 12–24-hour sleep and/or wakeful retention periods before subsequent memory testing. While these designs can control for condition differences due to initial behavioural accuracy increases following retrieval training [10–14], these methods may confound the effects of retrieval training with offline consolidation across subsequent sleep and wake periods [15]. These designs sometimes also confound the effects of retrieval training with the simultaneous restudy of other material, which may be subject to retrieval-induced forgetting [16]. Here we directly compare the separate impacts of sleep and retrieval training interventions across equivalent timescales to identify what transformations of memory representations are initially prioritised across both consolidation types.

Broad evidence exists for behavioural semanticization across memories and their transfer into neocortical networks across sleep [12,17–26] and retrieval training [7,27–35]. When memory representations are compared directly, increased representational overlap following sleep has been reported in the hippocampus and mPFC between objects paired with the same scene [36] and between stimuli with shared predictive structures [37]. For retrieval training, studies have also reported evidence for increasing representational similarity in the cortex and hippocampus within communities of items with similar predictive structures [38–40]. Overall, when memories are similar, neocortical integration across sleep and retrieval training typically leads to increased representational similarity between them (a phenomenon that we refer to as “representational merging”), driven by the maintenance of their shared conceptual details over the contextual details that initially differentiated them [41].

EEG offers a rich temporal approach that can measures the summation of local field potentials and the neural activity associated with memory representations. EEG phase patterns have been previously used to index memory reactivations, demonstrating sensitivity to category-specific and item-specific information [23,42–44]. Additionally, EEG sources can be reconstructed to provide spatial interpretability towards identifying item-specific and category-specific information [45]. Using representational similarity analysis with EEG, Liu et al [23] report that sleep is associated with increased category-level representation and reduced item-level representations. Applying a similar approach across retrieval training and sleep may offer further insights into aspects of memories that are undergoing representational changes to inform a comprehensive state-independent understanding of memory consolidation.

Although memory consolidation often supports greater memory accuracy and reduced interference [4,6,28,46], memory generalisation also results in a greater behavioural endorsement of critical lures (i.e., similar item lures) across sleep [47–49] and retrieval training [50–52]. When inter-item representational similarity between similar items is linked to behavioural outcomes, greater representational similarity is associated with an increased rate of false alarms and poorer memory fidelity after both sleep [53] and retrieval training [40,54]. Therefore, we would expect that representational similarity between similar items may relate to a greater likelihood of endorsing similar lures, following retrieval training and sleep.

Previous comparison between sleep and retrieval training that find similar evidence for generalisation have relied on different studies leveraging separate paradigms; yet, different paradigms can demonstrate opposing changes to memories [8,46,55]. Therefore, a comparison of sleep and retrieval training interventions in the same paradigm is important for drawing accurate conclusions on similarities in representational changes. Notably, Denis et al [51] measured the isolated impacts of retrieval training and sleep on memory generalisation across an audio story. They found that both sleep and retrieval training conditions led to more accurate veridical memory of story elements, as well as greater inference of non-presented, plausible material [51]. However, their design markedly did not include a restudy control group to rule out the effects of general memory practice, nor did they investigate a brain-based marker of representational changes to memories. Overall, the current state of evidence suggests that it has not been fully understood how the representational changes between memories compare across sleep and retrieval training.

The current study offers a direct comparison between retrieval-mediated and sleep-based consolidation’s ability to extract gist-like memory representations, linking fine-grained neural and behavioural changes. In four separate sessions, subjects learnt 104 different object-word pairings, where the objects across different pairs were similar lures to each other. Across a 120-min mid-afternoon period they then underwent either retrieval training practice (i.e., cued recall with feedback), restudy (i.e., seeing the pairs again), a nap opportunity, or wakeful rest. Subjects underwent immediate associative-recognition (pre-intervention) and delayed associative-recognition (post-intervention) testing of the object-word pairs as either old (original pairing) or new (rearranged pairing) with either similar- or different-object lures. EEG was measured during learning, immediate recognition, and delayed recognition phases to compare the reinstatement of memory representations of similar objects across time. Memory performance was measured at the item-level as binary accuracy to similar lures, and as session-level (i.e., per visit) recognition accuracy to similar-lures, different-lures, and original-pairings. Our results revealed an increase in phase similarity between similar objects (i.e., representational merging) across encoding, and further across the retrieval training intervention, but not sleep. Through this, we also investigated the times, frequencies, and sources where representational changes occurred the most across encoding and the interventions. Additionally, we observed a significantly lower improvement in similar-lure discrimination compared to original-pair identification across retrieval training. Lastly, although we predicted that representational merging would relate to reduced similar-lure discrimination, we found that greater representational merging predicted greater similar-lure discrimination performance across retrieval training.

## Results

The statistics summarising the demographic questionnaire results, subject-specific frequency band limits, and sleep characteristics during the sleep intervention are reported in S1 File. All subjects experienced some amount of NREM sleep during their nap opportunity. Only five subjects did not enter slow-wave sleep, and four subjects experienced rapid eye movement sleep. In this paradigm, subjects learnt object-word pairings, then underwent immediate and delayed associative-recognition testing either side of one of four 120-min interventions (retrieval training, restudy, sleep, or wake). A visualisation of the memory task can be found in Fig 1.

**Fig 1.**
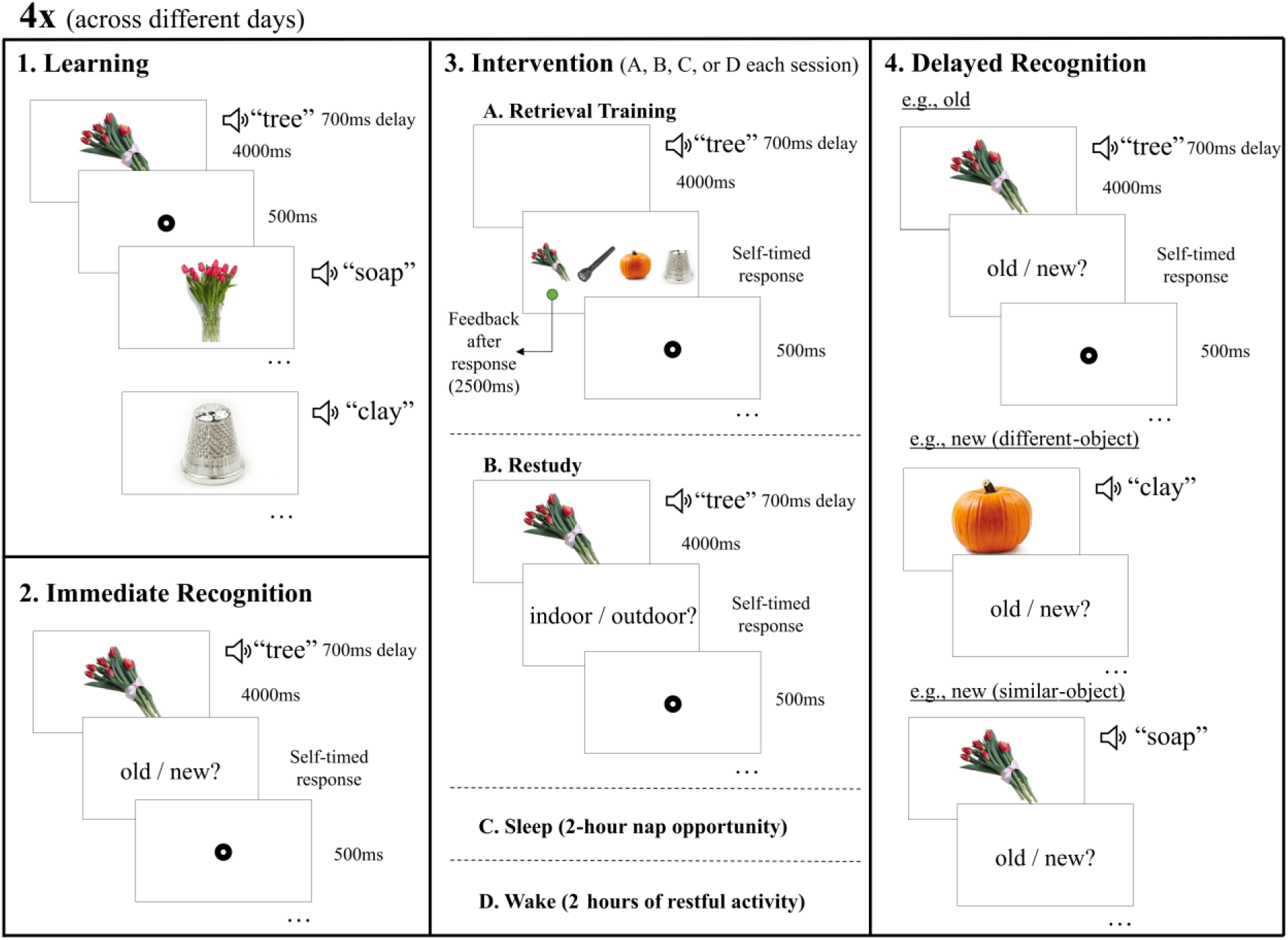
Memory task. 1. In the learning phase, subjects were exposed once to each of the object-word pairings. 2. Subjects then underwent immediate recognition testing, to distinguish between original (old) and rearranged (new) pairings, based on the pairings learnt during learning. An example of an old pairing is shown here. 3. Subjects underwent one of four intervention conditions. A. Involved retrieval training, where subjects heard each word, recalled the object, then picked it from distractors with feedback after. B. Subjects were exposed to all the pairs again and had to indicate if the object belonged indoors or outdoors. C. Subjects had a 120-min nap opportunity. D. Subjects engaged in light activities. 4. Delayed recognition involved the same procedure as immediate recognition, where subjects distinguished between old and new pairings of the objects and words. This includes an example of an old pairing, new pairing with a different object to the object paired with the original word, and a new pairing with a similar object to the object paired with the original word.

### Behavioural evidence for gist extraction across retrieval training

We first aimed to test if the change in recognition accuracy to each object lure type differed based on the intervention condition. A linear mixed-effects model found a significant interaction between intervention condition (i.e., retrieval training, restudy, sleep, and wake) and object type (i.e., similar-object lures, different-object lures, and original objects) to predict subjects’ change in recognition accuracy (scores compared between immediate and delayed recognition testing), *χ^2^*(6)=35.49, *p<*.001. The full model output is reported in S2 Table. This interaction effect is displayed in Fig 2. This figure demonstrates that retrieval training enhanced the identification of original pairings, but was the only condition to have significantly lower similar-object discrimination accuracy changes compared to the original-pair identification. Conversely, all accuracy types improved to a statistically similar degree across the sleep condition. Even though original object accuracy was numerically higher in the retrieval training condition, the testing effect did not reach significance as the 83% confidence intervals error bars overlapped between the conditions [56,57].

**Fig 2.**
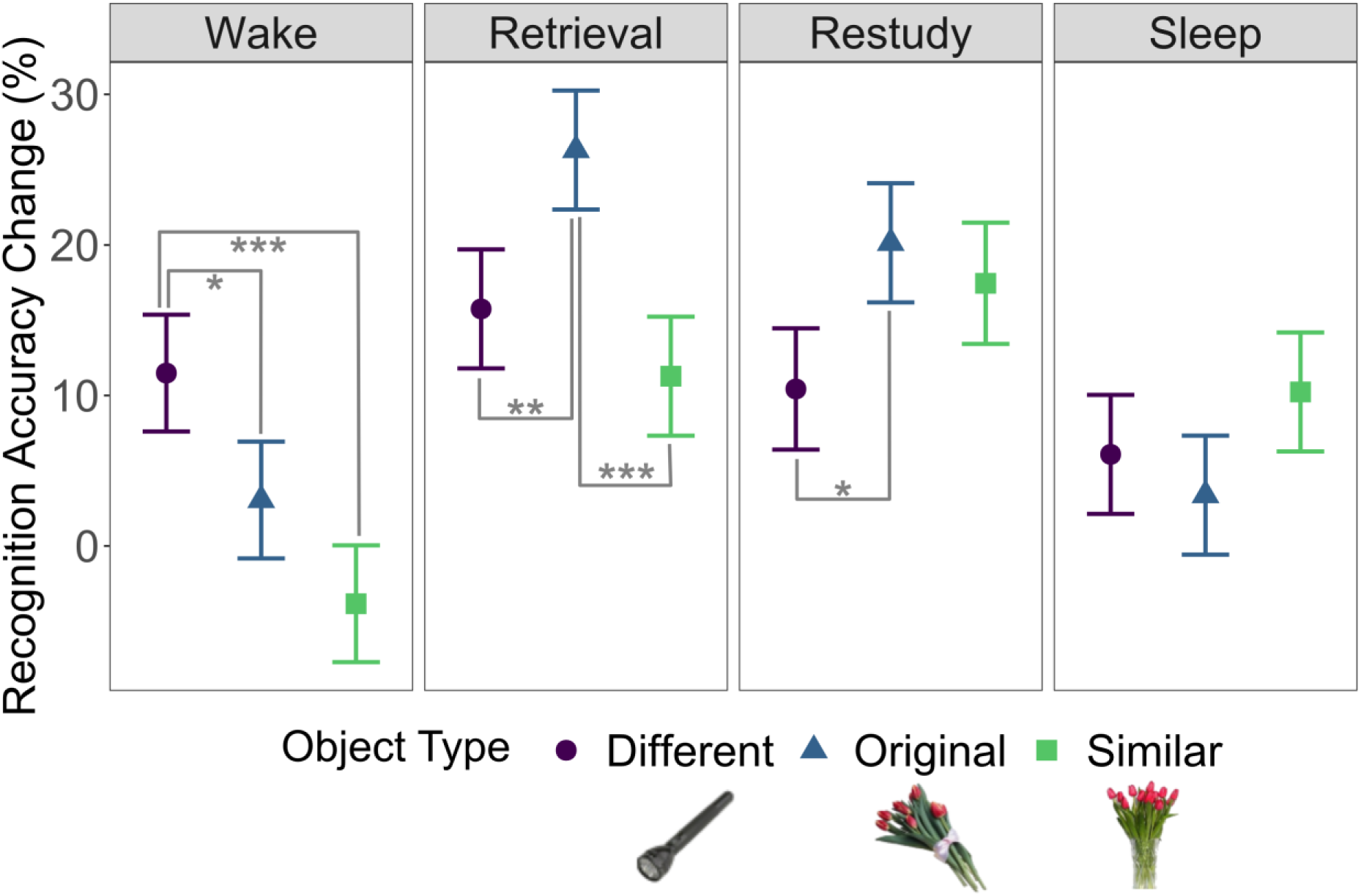
Condition differences for the change in recognition accuracy per object type. The y-axis represents the change in recognition accuracy percentage, from immediate to delayed recognition. Positive numbers indicate that there was a greater accuracy post-intervention, compared to pre-intervention. The object types include similar-object lures, different-object lures, and original object presentations to a paired word in the memory paradigm. The error bars represent the 83% confidence intervals, which reflects a 5% significance threshold for non-overlapping estimates. Significant contrasts are marked as: *denotes *p*<.05, \*\**p*<.01, and \*\*\**p*<.001.

### Encoding-driven representational merging of similar objects in theta-band phase

We next tested if similar objects became more similar in their phase angles across the initial encoding shift (i.e., representational merging from learning to immediate recognition). To test this, we first identified if object-specific information was maintained over the encoding shift through cluster-based permutation testing, which is reported in S3 File.

We further tested the direction of representational change across the encoding shift. This was done by computing representational change values between different object pairs and deriving a *p*-value as the number of grand average change-scores that were greater than the grand average change score between similar-object pairs. This analysis suggests that there was a significant shift towards greater phase similarity between similar-object pairs from learning to immediate recognition, significantly exceeding that of different objects (*p*=.002). This distribution is displayed in Fig 3a.

**Fig 3.**
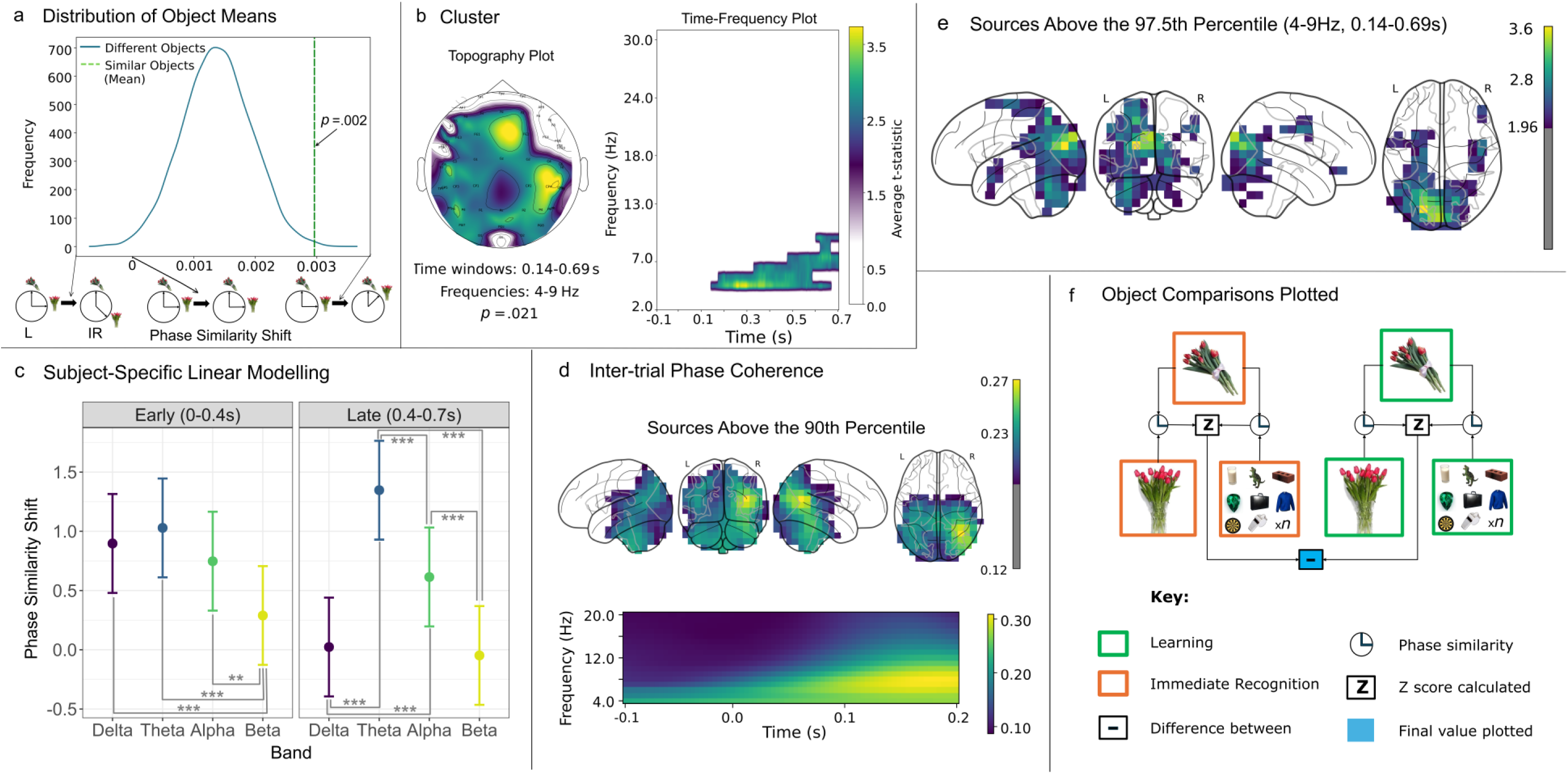
Representational merging from learning to immediate recognition. a) L = learning, IR = immediate recognition. The *p-*value highlighted in the distribution of object means refers to the number of different-object means that fall above the similar-object mean. These grand average values in the distribution are averaged across times, frequencies, and channels. The arrows and diagrams along the x-axis of the distribution graph illustrate the direction of change in phase similarity that occurs from learning to immediate recognition. Negative values indicate decreased phase similarity of similar objects at immediate recognition, positive values indicate increased phase similarity of similar objects at immediate recognition, and values closer to 0 indicate little change from learning to immediate recognition. b) The cluster of representational change z-scores for the encoding shift. The topography plot shows the channels contributing to the cluster, averaged over time and frequencies. The time-frequency plot shows the times and frequencies contributing to the significant cluster, averaged over channels. c) The interaction of frequency band and time window predicting representational change z-scores across the encoding shift. The frequency bands depicted have custom frequency ranges per subject, based on their individual alpha peak frequency. Negative values on the y-axis indicate decreased phase similarity of similar objects at immediate recognition, positive values indicate increased phase similarity of similar objects at immediate recognition, and values near 0 indicate little change from learning to immediate recognition. The facets represent the time windows. The error bars represent the 83% confidence intervals, which reflects a 5% significance threshold for non-overlapping estimates. Significant treatment contrasts are marked as: *denotes *p*<.05, \*\**p*<.01, and \*\*\**p*<.001. d) Source reconstruction of inter-trial phase coherence. L = left. R = right. The four glass brain plots on the left, depict the sources where the inter-trial phase coherence values between objects were above the 95th percentile of the data. The time-frequency plot shows the inter-trial phase coherence values averaged across occipital sources. e) Source reconstruction of representational change z-scores. These values are averaged across the times and frequencies depicted in the cluster in subfigure b. This plot depicts the sources that have the highest representational change z-scores across the encoding shift. The sources shown are above a z-score of 1.96, representing the 97.5th percentile. f) Illustration of the type of object comparisons that were performed for the representational change z-scores across the encoding shift (learning to immediate recognition) in this figure.

To pinpoint the times, frequencies, and channels contributing to this representational merging, a cluster-based permutation was conducted on the representational change z-scores (see Methods) across the encoding shift. These representational change z-scores represent the degree of change in similar-object over different-object representational similarity, across the encoding shift. This test identified one significant cluster representing a distribution where similar-object phase was more similar in the immediate recognition phase compared to learning (*p*=.021). The cluster involved frequencies of 4−9 Hz between 140−690 ms post-object onset, and was maximal over fronto-central channels (Fig 3b).

#### Linear mixed-effects modelling

Next, we tested if the representational merging significantly differed across broad time windows and subject-specific frequency bands to determine individual differences. For this analysis we adjusted subjects’ frequency band bounds based on their individual alpha frequency (IAF) values (see Methods). Therefore, we used a linear mixed-effects model to predict the representational change z-scores for the encoding shift as a function of subject-specific frequency bands (delta, theta, alpha, and beta) and time windows (early: 0−400 ms, late: 400−700 ms). This model revealed a significant interaction effect of time window and frequency band, *χ^2^*(3)=28.25, *p*<.001 on the representational change z-scores (see Table A in S4 File for the full model output), identifying that representational merging from learning to immediate recognition was most extreme in the late theta-band window. The time by band interaction effect is depicted in Fig 3c.

#### Source localisation

Lastly, we reconstructed the sources that contributed to the representational merging of similar objects across the encoding shift. To ensure that source reconstruction yielded plausible results with the filters we computed, we first localized the inter-trial phase coherence across all trials. Values above the 95th percentile within the distribution of data are plotted in Fig 3d, wherein the sources most involved in processing the initial onset of visual objects could be localized in occipital areas (note that, because the source reconstruction required that our data was referenced to and average reference, we also replicated the cluster-based permutations and distribution *p-*values for the average referenced sensor-level data. We report this analysis in Fig A and Fig B in S4 File).

We then reproduced the calculations of representational change z-scores (for the encoding shift) at the source-level to identify the contributing sources within the time and frequency windows of the sensor-level cluster (averaged at the source-level). Sources with representational change z-scores above the 97.5th percentile threshold are plotted onto a glass brain in Fig 3e and demonstrated a somewhat left-lateralised occipital cluster. A list of all the sources above the threshold, including their associated labels from the Harvard-Oxford anatomical atlas, are included in Table B in S4 File.

Overall, these results demonstrate that the phase similarity between similar objects increases from the first time of encoding to when they are tested. This representational merging occurred most prominently in the theta band and in occipital sources.

### Only the retrieval training intervention induces further representational merging of similar objects in the alpha-band

To address our hypothesis that the sleep and retrieval training conditions would undergo further representational merging, we conducted the same analyses across the intervention shift. To enable comparison, these analyses were performed separately for each intervention.

We first tested the broad direction of representational change across each of the four interventions. We derived a *p*-value as the number of grand averages of representational change scores for different object pairs that were greater than the grand average representational change between similar-objects. The resulting distribution *p-*values differed between the four conditions; only retrieval training (*p*=.005), but not restudy (*p*=.964), sleep (*p*=.545) or wake (*p*=.888), demonstrated a significant shift towards greater phase similarity (i.e., representational merging) for similar-objects beyond different-objects, from immediate to delayed recognition. The distributions for each condition are displayed in Fig 4a.

**Fig 4.**
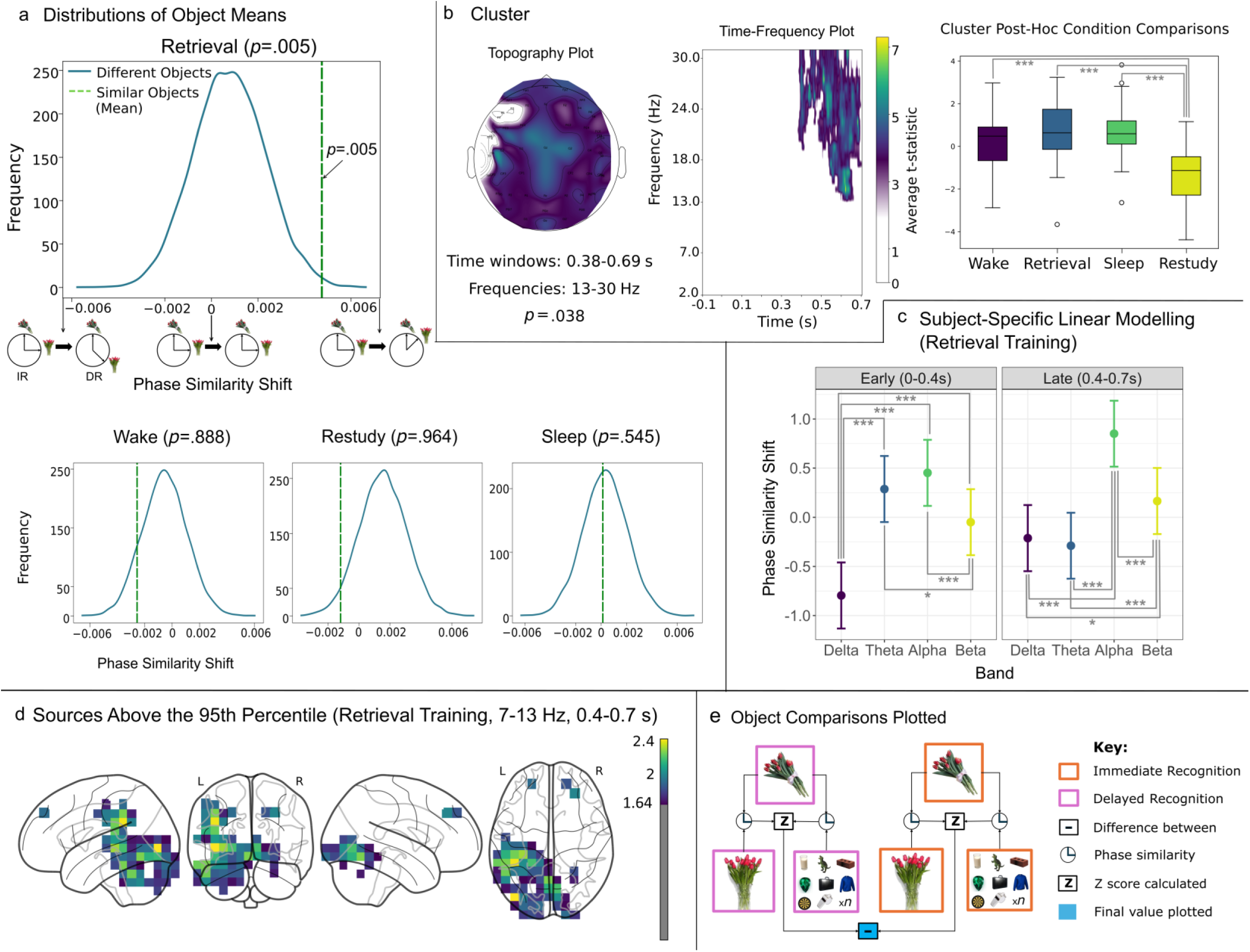
Representational merging from immediate to delayed recognition. IR = immediate recognition, DR = delayed recognition. a) The *p-* value highlighted in the distribution of object means refers to the number of different-object means that fall above the similar-object mean. The arrows and diagrams along the x-axis of the distribution graph depict the direction of change in phase similarity that occurs from immediate recognition to delayed recognition. Negative values indicate decreased phase similarity of similar objects at delayed recognition, positive values indicate increased phase similarity of similar objects at delayed recognition, and values closer to 0 indicate little change from immediate to delayed recognition. b) The cluster of representational change z-scores determined from the cluster-based permutation across time, frequencies, and topography, for the intervention shift. The topography plot shows the channels contributing to the cluster, averaged over time and frequencies. The time-frequency plot shows the times and frequencies contributing to the cluster, averaged over channels. The cluster post-hoc plot depicts the distribution of representational change z-scores across subjects (averaged per condition within the time, frequency, and channel limits of the cluster). Significant *t*-test comparisons are marked as: *denotes *p*<.05, \*\**p*<.01, and \*\*\**p*<.001. c) The interaction of frequency band and time window predicting representational change z-scores across the intervention shift in the retrieval training condition. The frequency bands depicted have custom frequency ranges per subject, based on their individual alpha peak. Negative values on the y-axis indicate decreased phase similarity of similar objects at delayed recognition, positive values indicate increased phase similarity of similar objects at delayed recognition, and values closer to 0 indicate little change from immediate to delayed recognition. The facets represent the time windows. The error bars represent the 83% confidence intervals, which reflects a 5% significance threshold for non-overlapping estimates. Significant treatment contrasts are marked as: *denotes *p*<.05, \*\**p*<.01, and \*\*\**p*<.001. d) L = left. R = right. The representational change z-scores at the source level, averaged across the late (400–700 ms) alpha band, which displayed the strongest representational merging (compare subfigure c). The plot depicts the sources with the highest representational change z-scores across the intervention shift, for the retrieval training condition only. The sources shown are above a z-score of 1.64, representing the 95th percentile. e) Illustration of the type of object comparisons performed for the representational change z-scores across the intervention shift (immediate to delayed recognition), that are plotted in this figure.

We next investigated the times, frequencies, and channels where representational merging differed between the four interventions. To this end, we conducted a cluster-based permutation test comparing representational change z-scores between conditions in a single test (representational change z-scores reflect the degree of change in similar-object representations over different-object representations, across the intervention shift). This test revealed one significant cluster wherein the change in representational similarity (i.e., greater similar-object similarity during delayed recognition over immediate recognition), had the greatest differences between the four intervention conditions (*p*=.038). This cluster encompassed frequencies between 13−30 Hz, between 380−690 ms post-object onset, and was distributed across a wide array of channels, visualised in Fig 4b. Post-hoc paired samples *t*-tests between the conditions revealed that this cluster was driven by the restudy condition, which displayed the least representational merging of all conditions, plotted in Fig 4b.

#### Linear mixed-effects modelling

To further elucidate the contributions of subject-specific frequency bands and time windows to representational merging, we next fitted a linear mixed-effects model for the retrieval training group. We modelled the average representational change z-scores were predicted by subject-specific frequency bands (delta, theta, alpha, and beta) and time windows (early: 0−400 ms, late: 400−700 ms) as fixed effects. The model detected a significant interaction effect of frequency band and time-window (*χ^2^*(3)=34.06, *p*<.001), on the representational change z-scores (see Table A in S5 File for the full model output). The interaction between time window and frequency band is depicted in Fig 4c. Treatment contrast coding revealed that the shift towards greater similar-object phase similarity in delayed over immediate recognition was strongest in the late (400−700 ms) alpha-band.

#### Source localisation

Finally, to reconstruct the sources underlying change in representational similarity of similar-objects across the retrieval training intervention, we calculated the representational change z-scores at the source-level (see Fig A in S5 File for the average reference sensor-level results). These source-level data were averaged across 400–700 ms in the alpha-band (as per identification in the linear-mixed effects model). Sources above the 95th percentile are plotted onto a glass brain in Fig 4d displaying a left-lateralised parieto-occipital distribution of sources, wherein representations of similar objects was more similar after the retrieval training intervention, compared to before the intervention. A list of all the sources above the threshold, and their anatomical labels, is included in Table B in S5 File.

In summary, our results suggest that representational merging of similar objects occurs only across the retrieval training condition, most prominently in the alpha band and in left-lateralised parieto-occipital sources. The representational similarity changes from immediate to delayed recognition also differed between intervention conditions in the late beta band, driven by differences to the restudy condition.

#### Item-level behavioural accuracy to similar-lure objects is predicted by representational merging in the retrieval training intervention

Finally, we tested if the representational merging of similar objects is related to subjects’ propensity to endorse similar lures. To this end, we tested if subjects’ item-level accuracy to similar lures after the retrieval training intervention could be predicted by that item’s representational similarity to its similar object before retrieval training (i.e., representational merging, see Methods for calculation details). We averaged the item-level representational change z-scores across the late time window (400−700 ms) and all alpha-band frequencies, as these were the time-frequency bins identified to most maximally undergo representational merging across retrieval training. We used these item-level representational change z-scores in a linear mixed-effects logistic regression to predict the binary accuracy to similar lures during delayed recognition (1=correct rejection, 0=false alarm). The overall model was significant, *χ^2^*(3)=4.78, *p=*.029, and correctly classified 82.20% of cases, based on the degree of representational merging of individual objects. The model indicated that for every unit of the item-level representational change z-score, subjects had a 3.49% greater likelihood of correctly identifying the lure as an incorrect pairing (OR= 1.03, 95%CI [0.01, 0.06]; depicted in Fig 5). In two additional control models, we predicted different-lure accuracy and original-pair identification accuracy from item-level representational change; however, no significant relationship was detected for either model (different-lure model: *χ^2^*(3)=1.25, *p=*.264, original-pair model: *χ^2^*(3)=0.01, *p=*.958), ruling out that representational change simply related to overall accuracy. See S6 File for the results of our models exploring session-level behavioural accuracy.

**Fig 5.**
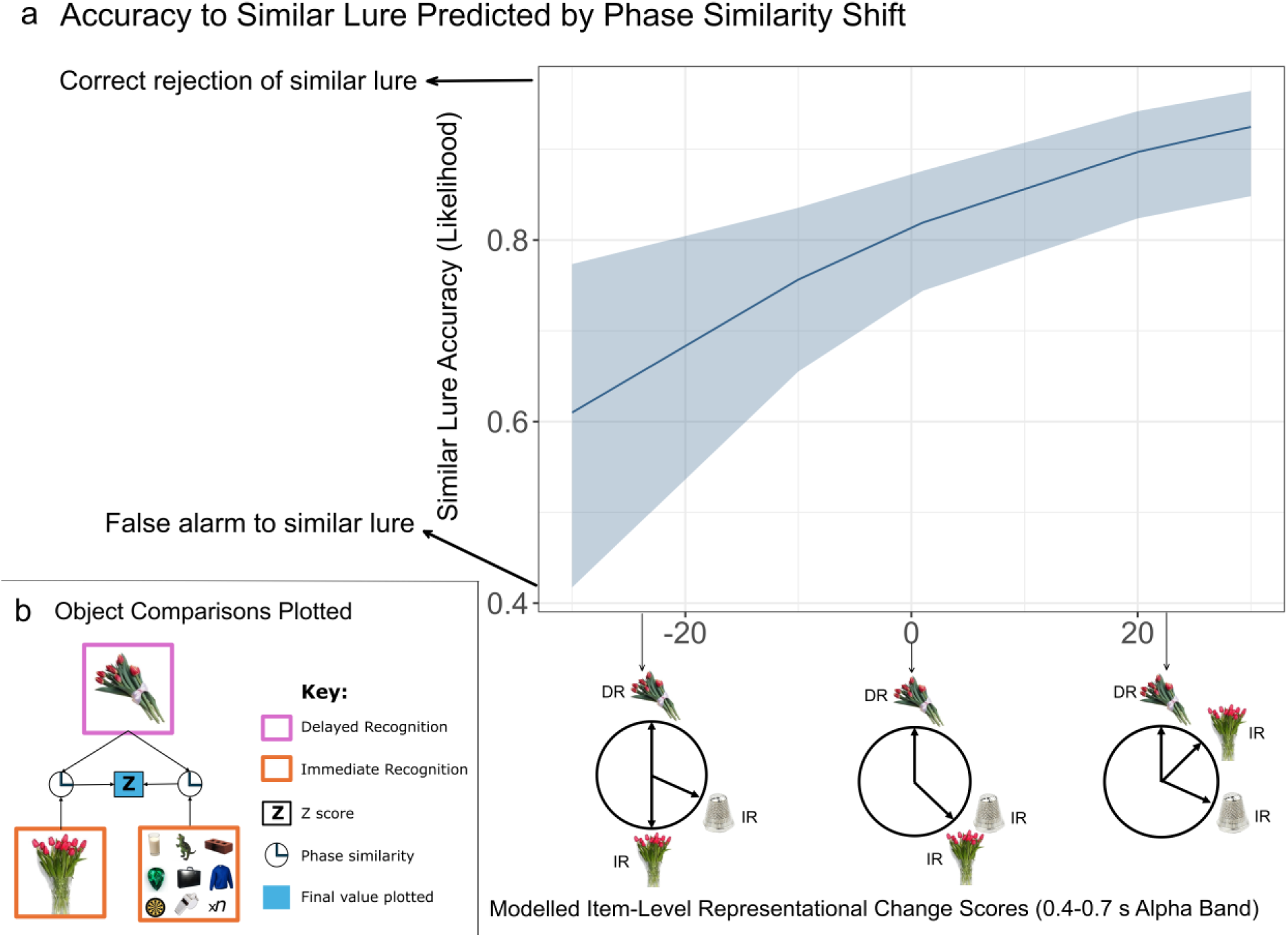
Item-level accuracy predicted by representational merging across retrieval training. IR = immediate recognition. DR = delayed recognition. a) The y-axis represents the modelled likelihood of subjects’ accuracy. Scores closer to zero indicate a greater likelihood of a false alarm to a similar lure, whereas numbers closer to one indicate a greater likelihood of a correct rejection of the similar lure. The x-axis represents the modelled modified item-level representational merging z-scores, which indicate the phase similarity of similar objects across time compared to different objects across time. The circular figures below the x-axis are a pictorial representation of an example relationship of phase similarities between similar and different objects that are captured by the negative, neutral, and positive values. Negative values indicate that an object at delayed recognition is more similar to different objects from immediate recognition than to its own similar pair-mate object from the immediate recognition phase (differentiation). Zero values indicated that an object at delayed recognition is as similar sleep condition. Subjects were seated in front of a dedicated computer and were fitted with an appropriately sized EEG cap. Subjects were then given details about the memory task they performed across the rest of the session.

## Discussion

In the current study, we tested how representations of similar objects change in EEG phase similarity across a daytime nap, retrieval training, wakefulness and restudy intervention. We further tested for behavioural differences in gist extraction between intervention conditions, and if similar-lure discrimination could be predicted by the degree of changes between the representations of similar objects in EEG phase patterns. We found evidence for reduced improvements to similar-lure discrimination, relative to the enhancement of original-pair identification, as well as the representational merging of similar objects that was confined to the retrieval training condition, and markedly absent after the sleep, wakefulness, or restudy intervention. Greater representational merging also predicted behavioural performance in the retrieval training condition, where a greater chance of correctly rejecting a similar lure was associated with greater representational merging of the similar object representations.

Only in the retrieval training condition did we observe a change in similar-lure discrimination that was significantly lower than the enhancements to original-pair identification. Across restudy, sleep, and wake interventions, the improvements to original-pair identification were not prioritised over a maintenance of the episodic fidelity used to discriminate similar lures. This aligns with previous research that retrieval training can induce a rapid extraction of gist that enhances overall memory accuracy, without an equivalent increase in episodic fidelity [50,51]. Across the sleep condition, we instead observed minor and insignificantly different changes between original-pair identification, different-object discrimination and similar-lure (i.e., episodic) discrimination. The sleep intervention not only seemed to protect episodic details from the decay experienced across wake (i.e., significantly lower similar-lure discrimination across wake compared to every other condition) [58], but it’s results may also reflect a selective maintenance of both the semantic and episodic information that were both integral to task goals [59]. This selective and protective nature of sleep differs from the retrieval training condition, which instead prioritised the enhancement of original-pair (i.e., semantic) identification through rapid gist extraction. This highlights a key difference in the types of memory transformations that occur across equivalent time periods of sleep and retrieval training interventions.

Our primary prediction was that representational merging would occur between similar objects across encoding, and importantly after both the retrieval training and sleep interventions, compared with restudy and wakefulness. By comparing the change in representational similarity of similar objects over different objects across encoding, we found evidence for a general trend towards increased representational similarity of similar objects from learning to immediate recognition. Across this encoding shift, increased similar-object representational similarity was greatest in the late (400−700 ms post-object onset) theta-band and was localised to the occipital lobe. This may reflect the initial incorporation of similar objects into similar neocortical networks across early encoding [60,61], and increased binding of similar bottom-up and coarse perceptual information between the pairs [62–64]. Crucially, this representational merging between similar objects only continued across the retrieval training condition, which was strongest in the late (400−700 ms) alpha-band and parietal-occipital areas. These time-frequency-source signatures may reflect the retrieval of shared semantic information and coarse perceptual similarities between objects [33,65–73] and thus the neocortical integration of similar pairs across consolidation [1,41]. However, we did not detect late alpha-band representational merging using the cluster-based permutation. This could suggest that representational merging across retrieval training occurred within subject-specific frequencies that were not similar enough between subjects to be detected in the cluster-based permutation.

The evidence for representational merging and gist extraction across retrieval training aligns with predictions that retrieval training is able to induce a rapid form of consolidation [1,6,8]. It is suggested that repeated reactivation drives competing memory representations to become more distinct from competitors [6,8]. However, the competitors used in most other studies have been stimuli introduced as interference with a task motivation to forget them or were new unencountered lures, opposed to task-relevant items that were also consolidated [20,74–77]. Instead, our study demonstrated that competitors are not always supressed, and representational merging can occur between competing memories when the task-demands require the maintenance of numerous competing items with similar features. Memory differentiation still could have occurred between competing memories, but was overshadowed by the overall trend towards representational merging that we observed, due to the constraints of scalp EEG measurements.

To investigate the functional implications of representational merging across retrieval training, we tested the prediction that greater representational merging would lead to a greater propensity to endorse similar lures. Our item-level model revealed a significant relationship between late (400−700 ms) alpha-band representational merging and accuracy at distinguishing similar lures. Previous studies found that the incorporation of similar objects into similar neocortical networks across consolidation, leads to poorer similar-lure discrimination due to gist extraction [40,53,54,78]. To our surprise, greater representational merging of two semantically-similar objects predicted an enhanced ability to behaviourally discriminate them. Follow-up analyses suggested that this relationship was not driven by a general association between representational changes and overall accuracy. This finding contrasts the session-level (i.e., average per visit) trends that we observed whereby episodic discrimination was significantly less enhanced across retrieval training compared to semantic discrimination, mirrored by representational merging in the EEG. This difference between session- and item-level trends adds nuance to our understanding that retrieval training universally provokes behavioural generalisation across memories through representational merging. In cases of greater representational merging across the retrieval-induced neocortical integration, episode-unique information was likely maintained. These transformations may allow for efficient storage and retrieval of unique episodic traces between similar memory representations to aid their discrimination [79,80]. This is supported by research where retrieval training is followed by sleep to further neocortically integrate memories. These studies demonstrated that the increase of category-specific neocortical representations does not come at the expense of episode-specific representations [33,81,82]. Our results may suggest that similar representational changes could be accomplished by some memories across retrieval training without subsequent sleep, but that this was overshadowed at the session-level by the majority of cases which undergo gist extraction. Future research should aim to determine was factors influence which memories undergo greater neocortical integration that supports greater episodic discrimination, such as encoding-related theta power [48].

Unexpectedly, we did not observe representational merging across the sleep intervention, despite previous studies predicting that representations of similar items becomes increasingly similar occurs across sleep [23,36,37]. However, studies have shown mixed results concerning gist extraction across single sleep episodes, where some paradigms do provoke gist extraction, and others do not, with differences in representational changes serving the unique goals of the paradigm [12,20,46,55,83]. This is why it was important to compare sleep and retrieval training in the same paradigm, where there are no inherent differences in paradigm-specific task goals. Therefore, whilst sleep-based consolidation effects are starkly task-specific, and sometimes need more time to come about, the effects of retrieval-mediated consolidation are consistently rapid and may be more robust across tasks. To support this claim, future studies should compare retrieval training and sleep across a wider variety of tasks with different goals, and link these representational changes to markers of consolidation that were not possible to detect in our paradigm (e.g., item-specific reactivations during consolidation interventions).

Our research begets various questions for future studies to address. Comparing consolidation using varying sleep lengths, stages, and microstructural events to retrieval training could illuminate the extent to which retrieval training is able to speed up memory transformations, and if any task-specific differences in consolidation occur. Also, investigating the longevity of these representational changes could reveal if retrieval training and sleep eventually lead to similar changes, and if these changes endure across time. Another route for exploration would be to experimentally vary the perceptual and semantic similarity between objects on a continuous scale, to see at what point these similarities between objects influences the degree of representational merging across consolidation. Lastly, our sample was comprised primarily of women, so future studies should investigate a more balanced sample to more confidently attribute these findings to the broader population.

Overall, the current study found evidence to suggest that representational merging occurred between similar-object memories across retrieval training, but not sleep. Additionally, we found that greater representational merging led to a greater likelihood of being able to distinguish between the merged memories. This revealed two interesting implications that are important for our understanding of memory consolidation. The first is that sleep and retrieval training are not equal in their consolidation of memories, when applied on the same time scale. The second is that whilst retrieval training may induce gist extraction on average, cases of greater neocortical integration can support episodic discrimination without the addition of sleep. Future research should continue to examine comparisons between sleep and retrieval training across different sessions, within the same paradigm, to further inform state-dependent and state-independent theories of memory consolidation.

## Materials and Methods

### Subjects

Thirty individuals (27 women, 1 non-binary, 2 men) aged 18–34 (*M*=22.17, *SD*=4.21) years participated in the study. Twenty-five completed all four conditions (retrieval training, restudy, sleep, and wake), one subject only completed retrieval, one subject completed restudy, one subject completed wake, and two subjects completed sleep and wake conditions. All collected data was used in the analyses. Subjects were right-handed, fluent English speakers, had normal or corrected vision, and self-reported no hearing or sleep problems. Subjects also reported upon interview and questionnaire no diagnosed psychiatric or sleep impairments, and were not taking any medications or recreational drugs in the last 6 months impacting EEG [84–86]. Subjects were recruited via flyers posted around the university campuses, social media, and an online subject recruitment system. Subjects who completed all four conditions received a $200 honorarium. The UniSA Human Research Ethics Committee granted ethics approval for this project (protocol no. 205130).

### Materials

#### Surveys

Sleep habits across the last month were screened using the Pittsburgh Sleep Quality Index (PSQI) [87]. The PSQI contains 19 items which are used to generate a total score where scores >5 indicate significant sleep difficulties, which was used as a cutoff for eligibility. The PSQI has demonstrated adequate internal consistency and high test-retest reliability [87–89].

The Morningness-Eveningness Questionnaire (MEQ) was used to screen out subjects with extreme circadian types [90]. The MEQ is a self-report questionnaire involving 19 multiple choice questions on sleep/wake habits. Subjects with scores ≤30 or ≥70 were excluded. The MEQ’s items have previously demonstrated a high internal consistency and high test-retest reliability [91,92].

Subjects’ handedness was screened using the Flinders Handedness Survey, where summed scores between −10 to −5 indicate left-handedness, −4 to +4 indicate mixed-handedness, and +5 to +10 indicate right-handedness [85]. Subjects with handedness-scores <5 were excluded.

The Karolinska Sleepiness Scale (KSS) was administered before the experiment began and again before the final testing to track subjects’ self-reported sleepiness [93]. The KSS comprised of a single-question, asking subjects to rate their current sleepiness on a scale from “1 – extremely alert” to “10 – extremely sleepy, cannot keep awake.”

Subjects’ age and gender were recorded via a custom questionnaire administered in *Qualtrics*. Additional screening questions asked about medication and drug use (e.g., “have you taken any recreational drugs in the last 6 months?”, see S7 Text for the full list of screening questions).

#### EEG recording

EEG was measured across each experimental phase, using a 64-channel passive electrode BrainCap with Ag/AgCl electrodes positioned according to the modified 10–20 system [94]. Electrooculographic activity was measured via two electrodes embedded in the cap, positioned approximately 1cm diagonally away from the outer canthus of each eye. Electromyographic activity was measured via three electrodes (two placed on the skin above each subglottic muscle, and one placed between the subglottic muscles), and electrocardiographic activity was measured by one electrode placed above the heart. Two BrainAmp amplifiers (DC system, sampling rate 500 Hz) were used in combination with BrainVision Recorder software for output recording. Impedances were kept below 10kΩ throughout all experimental phases.

#### Stimuli

We randomly sampled 416 object images (104 per condition) from sets 2–5 of the Mnemonic Similarity Task (MST) [95]. Each object image has another object image in the same set that is classed as a similar lure (e.g., a bouquet of flowers in a vase, versus a bouquet of flowers tied in a ribbon). To make sure the similar object lures were discriminable from each other above chance level, we only included MST images where the similar-object lure was endorsed less than chance level (i.e., ≤33.3%) in the MST’s accompanying dataset.

Word stimuli were taken from an updated version of the Affective Norms for English Words [96]. This dataset contains 13915 English words, optimised for use in paired associate tasks, rated on frequency from 0–1000 and 1–9 on valence, arousal, and dominance. This study randomly selected 416 monosyllabic nouns (104 per condition), with valence and arousal ratings between 3.5–6.5. These words were recorded, using the voice of a native English speaker with an Australian accent, for their auditory presentation. These recordings were approximately 500–1000 ms long, each. Words were presented at approximately 70 dB.

### Procedure

Eligible subjects were screened using the PSQI, MEQ, and Flinders Handedness Survey. Subjects then participated in four conditions, each of 4.5 hr (12:00–16:30; see Fig 6), separated by at least 48 hr. The night before each condition, subjects were asked to wake up 1 hr earlier than usual to increase the likelihood of an afternoon nap entering NREM sleep, in the sleep condition. Subjects were seated in front of a dedicated computer and were fitted with an appropriately sized EEG cap. Subjects were then given details about the memory task they performed across the rest of the session.

**Fig 6.**
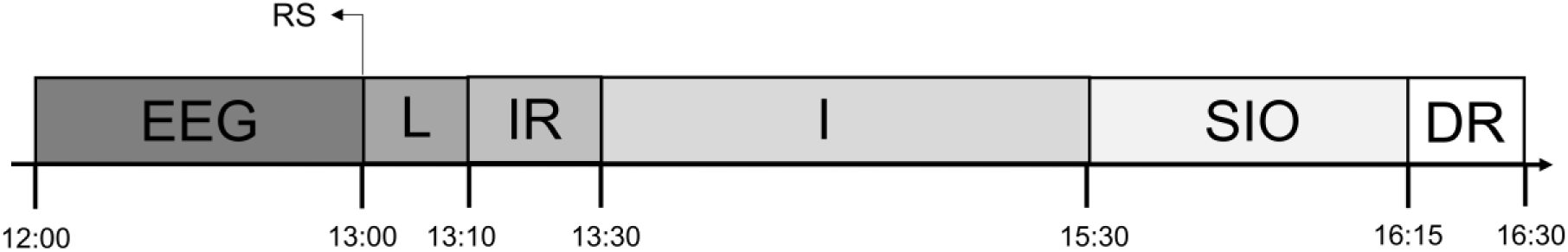
Procedure timeline. EEG = Setup EEG, RS = Resting State, L = Learning, IR = Immediate Recognition, I = Intervention (retrieval training, restudy, sleep, or wake), SIO = Sleep Inertia Offset / Retention Period, DR = Delayed Recognition. The memory task was based on previous protocols which have employed retrieval training and sleep [44,82,91]. *Open Sesame* v.3.3.10 software was used to create and run each phase of the memory task [97]. The memory task began with a learning phase, then immediate recognition, one of four intervention conditions (retrieval training, restudy, sleep, or wake), and a final delayed recognition. The intervention conditions across sessions were experienced in a pre-determined, counterbalanced order. This counterbalancing was random, except that the subjects always completed one wakeful condition before the sleep condition, to increase their familiarity with the environment and therefore increase their likelihood of napping. The paired associates used in the memory task consisted of randomly paired MST object pictures and auditory words. All subjects were presented with the same object-word pairs, in a randomised order for each condition, by subject. A different set of object-word pairs were used in each of the four conditions. For the object in each object-word pair, there was an object in a different object-word pair that was its similar object from the MST. Highly related object-word pairs were manually identified and reshuffled. The following sections detail each experimental phase of the memory task.

The memory task was based on previous protocols which have employed retrieval training and sleep [44,82,91]. *Open Sesame* v.3.3.10 software was used to create and run each phase of the memory task [97]. The memory task began with a learning phase, then immediate recognition, one of four intervention conditions (retrieval training, restudy, sleep, or wake), and a final delayed recognition. The intervention conditions across sessions were experienced in a pre-determined, counterbalanced order. This counterbalancing was random, except that the subjects always completed one wakeful condition before the sleep condition, to increase their familiarity with the environment and therefore increase their likelihood of napping. The paired associates used in the memory task consisted of randomly paired MST object pictures and auditory words. All subjects were presented with the same object-word pairs, in a randomised order for each condition, by subject. A different set of object-word pairs were used in each of the four conditions. For the object in each object-word pair, there was an object in a different object-word pair that was its similar object from the MST. Highly related object-word pairs were manually identified and reshuffled. The following sections detail each experimental phase of the memory task.

#### Learning

In each session, subjects learnt 104 object-word pairs. This involved viewing the image of an object on a white screen for 4000 ms and a sound of the paired auditory word was played 700 ms after the object onset. The images were displayed in colour, 400 pixels wide, in the centre of the screen. After each object was displayed, a fixation dot (8 pixels wide with a 2-pixel wide centre hole) was presented for 500 ms, before the next object was shown. Each pair was presented once in a randomised order.

#### Immediate recognition

Immediately following learning, recognition memory of the pairs was tested by exposing subjects to each object and word again in the same sequencing as the learning phase (i.e., shown an object, hear the paired word), with the pairs presented in a randomised order. This was done using an old/new paradigm, where 72 object-word pairs were presented in their correct pairings (old pairs), and 32 in a rearranged pairing (new pairs). To create the rearranged pairings, pairs were shuffled by substituting the object in each pair with another object of a different pair (16 similar objects, 16 different objects). After each pair was presented, a new white screen displayed the prompt “old or new?” Subjects responded via keypress, based on whether they believed that the pairing was presented as it was originally learnt (old / “Q”), or if it was rearranged (new / “P”). Before starting the task, subjects were encouraged to respond quickly, but optimize accuracy over speed. After they responded to the prompt, a fixation dot appeared on the screen for 500 ms before the next object-word pair was shown. No feedback was given to subjects on recognition accuracy performance. To avoid introducing interference effects with the original encoding episodes, the 32 rearranged (new) pairs were not used in subsequent phases.

#### Interventions

For each of their four visits, subjects completed one of the four interventions, depending on the condition order they were assigned. Each intervention period took 120 min and EEG was recorded throughout, apart from one exception, the wake condition. After the intervention, all conditions included a 45-min retention period, which also served as a sleep inertia offset period for the nap condition. During this break, subjects remained in the laboratory and engaged in light activities (e.g., reading or scrolling on social media), simulating how they typically spend their free time. The KSS was completed at the end of the intervention. The four interventions are outlined below.

Subjects completed six rounds of retrieval training practice with the initially learnt object-word pairs. The 72 object-word pairs were randomly divided into two equal length lists, with each list being presented three times. Within each round, the order of presentation for the object-word pairs was randomised per subject. Practice round took 10−12 min. Subjects had 8–10-min breaks between each round so that the practicing was spaced out over the 120-min period. The retrieval training practice had subjects hear the word of each pair, recall the paired object from their memory, then pick the object that was originally paired with the word from distractors. Subjects were presented a blank white screen and heard the word of one of the object-word pairs that were previously learnt. Subjects were instructed to imagine the paired object on the screen, when they heard the word. After 4000 ms, four object images that were part of previously learnt pairs were displayed 120 pixels wide, equally spaced across the width of the screen. Of the four objects, one of them was the original object that the word was paired with, while the other three were objects from other pairs. The distractor options were pre- determined and consistent across all subjects, but they were different each of the three times that a word was practiced. For every single pair that was practiced, exactly one round included the original object’s similar MST object as one of the distractors. Subjects were instructed to choose the object that they believed they previously learnt as paired with the word they just heard. Each distractor object had a black letter displayed above it which indicated the corresponding response key (Q, E, I, or P) required to select the object below it. 500 ms after their response, a green dot (8 pixels wide with a 2-pixel wide centre hole) was displayed below the correct object for 2000 ms to provide feedback, regardless of their accuracy. Subjects were instructed to use this feedback to learn the pairings and improve their memory. After the feedback was presented, a fixation dot was shown in the middle of the screen for 500 ms, before the next word was played on a blank screen. For the breaks between each round, subjects engaged in light activities.

In the restudy condition, subjects followed the same six-round study-break structure as the retrieval training intervention, practicing half (36) the 72 pairs at a time, three times each. Overall, restudy practice involved being exposed to both constituents of each word pair again. For each object-word pair, subjects were presented the object and word exactly as was done during learning, in a subject-specific randomised order. An object image from each pair was presented in the centre of the screen for 4000 ms, followed by the originally paired word presented auditorily 700 ms later. To maintain subjects’ attention, after each pair was presented, a screen prompted subjects to decide if the object of that pair would be commonly found indoors or outdoors. This prompt was displayed in the centre of a white screen, with the words “Indoors / Outdoors?” with the letter “Q” and underneath the word “Indoors,” and the letter “P” underneath “Outdoors.” These letters underneath the question depicted the keypress required to select the corresponding option. Subjects were instructed to treat this interactivity task as secondary to their practicing of the pairs. After subjects responded, a fixation dot appeared for 500 ms before the next object-word pair was presented.

Subjects were given a 120-min nap opportunity for the sleep condition. Subjects slept in a quiet, darkened bedroom, and to promote sleep white noise was played at approximately 37 dB during the entire nap opportunity. The researcher remained in the adjacent room, monitoring the EEG signal for signs of sleep and any disturbances to the electrode signals.

In the wake condition, subjects were supervised while they engaged in light activities (e.g., reading a book or watching a movie) for 120 min, to replicate their typical use of free time. During this time, subjects’ EEG was not recorded, but they remained in the laboratory with the EEG cap on.

#### Delayed recognition

Comparable to the immediate recognition phase, subjects were presented with the same old/new task, but with a new assignment of pairs as either old or new. Subjects were exposed to the 72 object-word pairs they practiced, which were presented using the same sequencing as the immediate recognition phase. For half (36) of the 72 pairs, the word of each pair was presented with its originally paired object from the learning phase. The other half (36) of the object-word pairs were novel (new) pairings. Half of the novel pairings were similar-object lures, whereas the other half were different-object lures. In the exact same structure as the immediate recognition phase, subjects were prompted, by the screen following the object, to indicate if the pairing was old or new, receiving no feedback on their accuracy. After they answered, a fixation dot appeared for 500 ms before the next pair was presented. Whilst the order of presentation of pairs was randomised for each subject, all shuffled pairings and assignments to old or new categories were pre-determined to remain consistent across all subjects. After the delayed recognition test, subjects had the EEG cap removed, and then left the laboratory. Subjects received their payment once they had finished their final session.

### Data analysis

#### EEG preprocessing

The EEG was pre-processed using *MNE-python* v1.7.0 [98]. Activity from the learning, immediate recognition, and delayed recognition phases was first resampled to 250 Hz, then re-referenced to mathematically linked mastoid electrodes. The data were then filtered with a zero-phase finite impulse response filter using a Hamming window with a transition bandwidth of 1–40 Hz. An Independent Component Analysis was performed to correct electrooculographic (EOG) artifacts by rejecting components which correlated the most strongly with EOG events (via the *create_eog_epochs* function in *MNE-Python*). Epochs were then created, starting −2500 ms before stimulus (object image) onset events and ending 6000 ms after stimulus onset (these large epochs were required to sample sufficient cycles in the time-frequency decomposition). Spectrograms were later cropped to −100−700 ms around the object onset. Finally, the *AutoReject* package [99] was used to reject bad epochs and repair bad channels via topographic interpolation. Sleep scoring of EEG data was performed as per set procedures [100]. Sleep stages 3 and 4 were combined for a joint slow-wave sleep measure.

Subjects’ IAF values were used to create subject-specific frequency band limits for later statistical modelling. IAF was calculated on the EEG 2-min eyes-closed recording, during which alpha activity is typically most prominent [101]. This was represented by the centre of gravity, a weighted average of the power within the alpha band (∼7–13 Hz) that circumvents problems with identifying a dominant alpha peak. The alpha centre of gravity was calculated in MNE Python, using methods from Corcoran et al [102]. This IAF value was then used to define subject-specific frequency band limits using the golden mean-based algorithm described in Pletzer et al [103].

#### EEG phase similarity

EEG phase similarity was calculated between different combinations of object types, across learning, immediate recognition, and delayed recognition. This was done within each session (i.e., one visit), at each channel, sampling point, and 1 Hz frequency step. Specifically, time-frequency representations were computed as complex values (spectral coefficients) for each epoch, using a Morlet wavelet transformation for 5 cycles per 1 Hz frequency step between 2−30 Hz, per 5 ms step. To compare the phase similarity between two objects, the complex spectral coefficients were first normalized to unit length and then averaged between trials corresponding to two objects. The length of this average vector was then taken as a measure of phase similarity. Comparisons between objects were done within each subject and condition, and at all 56 EEG channels, 29 frequencies and 200 time samples within the −100−700 ms epochs. The two methods of computations involving these EEG phase similarity calculations, are described below.

First, to initially test for the representation of object-specific information in EEG phase, phase similarity was calculated between trials where the same object was presented during learning and immediate recognition. Phase similarity was then compared between the object’s trial in learning and trials from a randomly selected different object in the immediate recognition period. The difference in phase similarity between the same-object and different-object comparisons was taken to capture object-specific phase (i.e., same-different comparison). Next, phase similarity was calculated between each object in learning and its similar object (the lure object based on the MST) during immediate recognition. The difference between same-object comparisons and similar-object comparisons was taken to capture the object-specific phase beyond the conceptual likeness it shares with the similar object (i.e., same vs. similar object comparison).

Second, to test representational change across the encoding and intervention shifts, phase similarities between similar-object pairings within learning, immediate recognition, and delayed recognition were standardised to a distribution of phase similarities from different-object pairings within the same experimental phase. These standardised values are referred to as z-scores hereafter. Difference scores were taken between these z-scores for the learning to immediate recognition phases (encoding shift), and for the immediate and delayed recognition phases (intervention shift). For the encoding shift, the data from every intervention condition was taken together to form the random different-object distributions, whereas for the intervention shift, separate distributions were calculated per intervention condition.

Specifically, the following steps were done to create a sample-level distribution of different-object phase similarities, per experimental phase. First, we only kept the object epochs which had a matching similar-object epoch, in each session (these differed per session due to different epoch rejection rates). We then calculated the minimum number of epochs across sessions (*n*=34 for the encoding shift, *n*=14 for the intervention shift) as this would form the minimum possible number of different-object comparisons that we could uniformly perform across all sessions. Next, we calculated the phase similarity between each object and the minimum possible number of random different objects. We then calculated average different-object values per subject, equal to the minimum possible number, by averaging across the objects. This resulted in 34 different-object average values across the encoding shift (*n*=14 across the intervention shift), per session. To create the sample-level distribution of different-object average values, one of these values was randomly chosen per session (i.e., 105 values were chosen), to calculate an average value across this subset. This cross-subject averaging was repeated 10,000 times to manually create a distribution of 10,000 sample-level different-object phase similarities (referred to hereafter as the baseline different-object distribution), from which the mean and standard deviation were taken ***[104]***. An average similar-object phase similarity value was also calculated, per session.

Representational similarity z-scores were then calculated per session, using the baseline different-object distribution mean and standard deviation, and the session-level similar-object phase similarity values. Lastly, a final representational change z-score was calculated by subtracting the difference between the representational similarity z-scores, from learning to immediate recognition (encoding shift), and from immediate to delayed recognition (intervention shift). Taking difference scores between these timepoints isolates the representational changes that occur across encoding versus the interventions. Fig 7a displays a visualisation of the type of object comparisons made across the encoding shift, as an example. Importantly, these two sets of representational change z-scores represent the degree of change in similar-object phase similarity, above that of different-objects, across initial encoding and the separate interventions. Positive values of the representational change z-scores indicate that the phase of similar-objects becomes more similar across the period (representational merging), whereas negative values represent similar-object phases becoming less similar across the period (representational differentiation).

**Fig 7.**
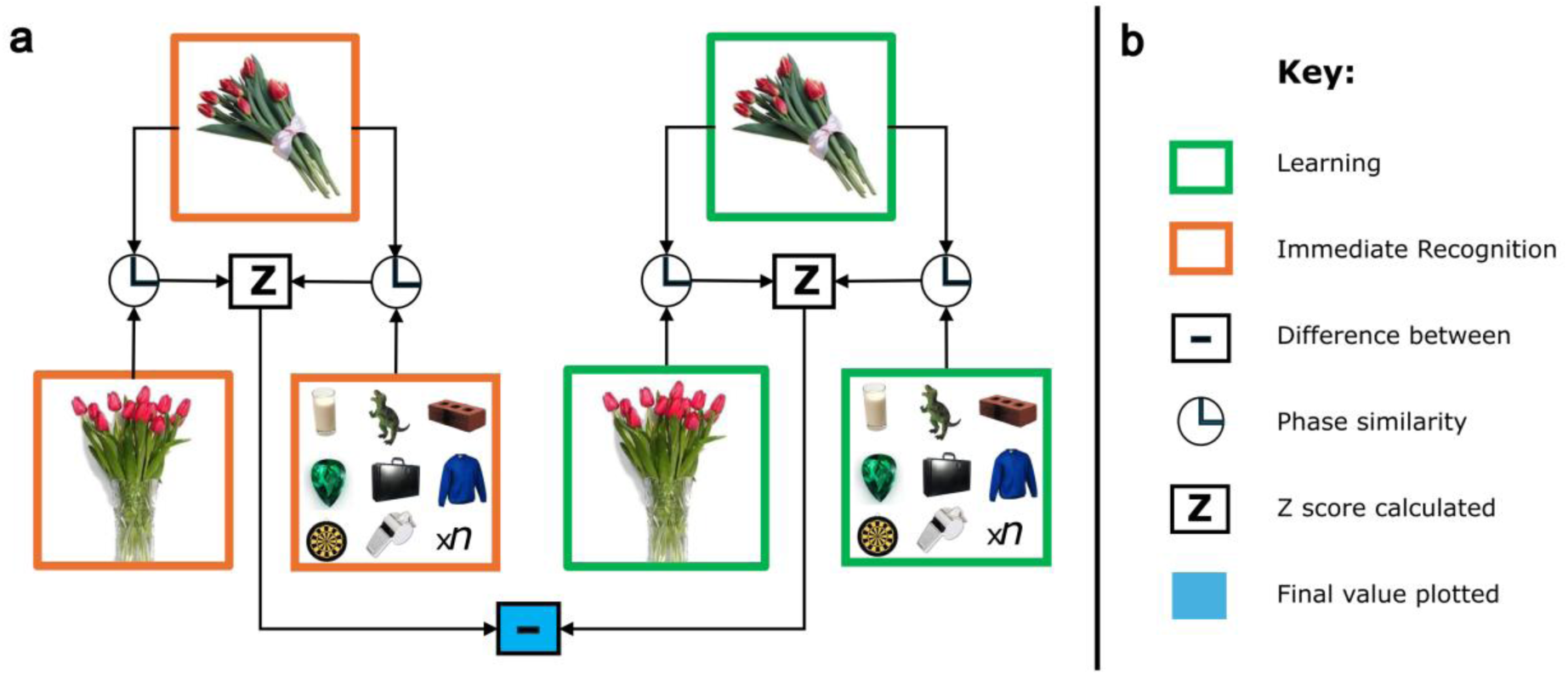
Illustration of phase similarity object comparisons. a) Object comparison across the encoding shift. Phase similarity was computed between an object in learning, and its similar object also in learning. A distribution was created of cross-subject means of phase similarities between an object in learning and *n* (*n*=34 for the encoding shift, *n*=14 for the intervention shift) random different objects in learning. This distribution was used to standardise the similar-object phase similarities and the entire process was repeated for immediate recognition. Finally, a difference score was created between learning and immediate recognition z-scores. This process was also repeated across the intervention shift between immediate and delayed recognition for each condition. The final scores were taken to represent the change in representational similarity across the encoding and intervention periods. b) Contains the key for subfigure a.

#### Statistical analyses

The following analyses were used to test our hypotheses. Firstly, we tested for differences in recognition accuracy to original object pairings and their lures, within the four intervention conditions. Secondly, we looked at four different aspects of representational change across the encoding shift (learning to immediate recognition) and the intervention shift (immediate to delayed recognition). To this end, we estimated the grand average trend of representational change across the encoding and intervention shifts per condition by calculating the proportion of different-object phase similarity means in a distribution that were greater than the similar-object mean. To identify the fine-grained specifics of these trends, we then computed four cluster-based permutations to identify times, frequencies, and channels associated with object-specific representations, and representational changes across the encoding and intervention shifts. We also conducted linear mixed modelling to identify the broader time and individual differences in subject-specific frequency windows in which this representational change occurred, per condition. We then estimated the anatomical sources associated with the representational changes occurring across the encoding and intervention shifts. Thirdly, to test if these representational changes impact the propensity to endorse similar lures, we linked these representational changes to session-level and item-level behavioural accuracy. Each step is described in detail in the following sections.

#### Behaviour modelling

To represent the endorsement of different object types at immediate and delayed recognition tests, we calculated the session-level (i.e., per visit) percentage recognition accuracy, per object type (i.e., similar-object lures, different-object lures, and original objects). For example, similar-object recognition accuracy reflects the percentage of correct rejections when exposed to similar-object lures. We then calculated change scores by subtracting immediate recognition accuracy from delayed recognition accuracy scores. Positive recognition accuracy change scores indicate improvements in recognition accuracy from immediate to delayed recognition tests.

To determine if the endorsement of each object type differed across intervention conditions, we conducted a linear mixed-effects model in *R* v.4.3.0 [105] using the *lme4* package [106]. This model used recognition accuracy change scores as the outcome, with intervention condition (retrieval training, restudy, sleep, or wake) and object type as fixed effects, with subject ID as a random effect. In all linear modelling conducted, outliers were removed before each model was conducted. Values were determined to be outliers if they were more than 1.5 times the interquartile range above quartile 3, or below quartile 1. To plot the significant main and/or interaction effects from each model, the modelled effects were extracted using the *effects* package [107]. To identify significant differences in recognition accuracy between intervention conditions and object types, treatment contrast coding was applied between object types within each intervention condition. In these plots, the error bars depicted the 83% confidence interval, which reflects a 5% significance threshold for non-overlapping estimates [56,57].

#### Representational change analyses

Firstly, we assessed the grand average direction of change in representational similarity of similar objects across the encoding shift (comparing learning to immediate recognition) and the intervention shift (comparing immediate to delayed recognition). To this end, we computed distribution *p*-values from the number of phase similarity values in a baseline different-object distribution that fell above the sample-level mean of similar-object phase similarities. These distribution *p*-values were calculated separately per condition for the intervention shift, but included all conditions for the encoding shift. To calculate these *p*-values, difference scores were computed between the baseline different-object distribution values of the two time points, per shift (between learning and immediate recognition for the encoding shift, and between immediate and delayed recognition for each condition in the intervention shift). The distribution *p*-values were averaged across channels, frequencies, and time, to produce 10,000 grand average values reflecting a distribution of different-object phase similarity values across time. Difference scores were then calculated between the grand average similar-object phase similarity values, per shift (i.e., encoding shift, and intervention shift per condition). Finally, the number of grand average different-object values that fell above the grand average similar-object value was calculated, to reflect the distribution *p-*value. Distribution *p-*values <.025 (two-tailed) conveyed that the similar objects had undergone representational merging across the shift that was statistically greater than that experienced between different objects. Conversely, distribution *p-*values >.975 (two-tailed) reflected that the similar objects had undergone greater representational differentiation across the shift.

Next, we aimed to identify the channels, frequencies, and times contributing to object-specific representations from learning to immediate recognition, as a sanity check that object-specific information was maintained over time in the EEG phase. To do this, we ran two cluster-based permutation tests (using *MNE-Python*’s *mne.stats.spatio_temporal_cluster_test* function), one with the EEG phase similarity for same-different object comparisons, and one for the same-similar object comparison data. Both cluster-based permutation tests used a two-tailed one-sample *t*-test test statistic.

To test for channels, frequencies, and times contributing representational changes across the encoding shift and intervention shifts, we ran another two cluster-based permutation tests. For the cluster-based permutation test investigating the encoding shift, we input the representational change z-scores for the encoding shift (i.e., change in similar-object phase similarity, above that of different-objects, from learning to immediate recognition), using a one-tailed one-sample *t*-test test statistic. For the intervention shift, a single test compared the representational change z-scores (i.e., change in similar-object phase similarity, above that of different-objects, from immediate to delayed recognition) between the four intervention conditions, using a one-way ANOVA test statistic. Post-hoc paired-samples *t*-tests were conducted on the representational change z-scores between all conditions for the intervention shift cluster test. As six *t*-tests were conducted, a Bonferroni correction for multiple comparisons was applied to adjust the alpha level (i.e., *p*=.008).

For all cluster-based permutation tests, adjacency was defined between neighbouring frequencies, times, and channels, and the test was computed with 1000 permutations. The cluster forming thresholds for all permutations were calculated according to their respective test types, using *t.ppf* and *f.ppf* from the *SciPy* package ***[108]***. Clusters found by the test that were *p*<.05 were considered significant.

Finally, to test whether changes in representational similarity across the encoding shift and the intervention shift differed across subject-specific frequency bands and time windows, linear mixed-effects modelling was conducted. To model changes across the encoding shift, the representational change z-scores were used as the outcome, averaged across time window bins (early: 0−400 ms, late: 400−700 ms) and frequency band bins (delta, theta, alpha, and beta, adjusted based on the subjects’ IAFs). The fixed effects were the frequency band and time window bins, with subject IDs and EEG channels as random effects. A similar model structure was created for the intervention shift, except that separate models were run per intervention condition to prevent multicollinearity between intervention condition and representational change z-scores (subsequently, subject ID was not needed as a random effect). Additionally, only the intervention conditions whose distribution *p-*value was <.025 (i.e., where there was evidence for representational merging), were modelled at this step. Treatment contrast coding was applied to this model to identify significant differences between representational change z-scores of each frequency band, within the same time-window.

#### Source localisation

To identify the sources underlying our observed sensor-level effects, we computed source localisations. Most of this data analysis remained the same as the sensor-level pipeline described above to calculate the representational change z-scores, but with a few key changes described hereafter. We re-referenced our data to an average reference to ensure the accuracy of the forward model. Epochs were not rejected anew for this pipeline; instead, to limit differences between the original and current pipelines, we rejected the same epochs identified in the original pipeline. We then replicated the same-different and same-similar cluster-based permutation tests with the sensor-level average referenced data, to see if object-specific information was still maintained across the encoding shift.

Source localisation was then calculated using the newly pre-processed source-level epochs in learning, immediate recognition, and delayed recognition trials. An MRI template brain scan from FreeSurfer [109] was used across all subjects. The pre-calculated boundary element model and transformation within the FreeSurfer dataset were used. A volume source space was sampled with 10mm spacing and a 4.75mm minimum distance from the bounding surface. The forward solution was computed for use across all subjects, as the MRI scan and electrode coordinates did not differ per-subject, resulting in 1844 sources. The inverse model was then calculated via linearly constrained minimum variance (LCMV) beamforming. Session-specific LCMV filters were calculated based on the data covariance across the post-stimulus section (0−700 ms) of the epochs with unit-noise gain and a regularisation of 5% (which accounted for rank deficiency of the data). Source orientation was fixed to the orientation maximizing power.

After applying the inverse solution, we calculated source inter-trial phase coherence at 8 Hz, between −100 ms pre-object-onset to 200 ms post-object-onset, across all trials during the immediate recognition experimental phase. This contrast served to establish the LCMV filter quality and to localise phase alignment due to stimulus perception. We then identified the sources contributing the most to the representational similarity changes by replicating analyses and statistical tests from the sensor level analysis, in source space. The calculation of the representational change z-scores across the encoding and intervention shifts were done in an identical pipeline as was done at the sensor level; however, for the source data the data were bootstrapped only 100 times, due to processing limitations of the considerably larger source-level data. We visualized the source-level analysis for the conditions and time-frequency bins identified in prior sensor-level modelling, to illustrate most plausible sources of known effects. The plotting threshold for the representational change z-scores in the intervention shift source map was set to the 95th percentile (1.645). For the encoding shift, the threshold for plotting was instead set to a more conservative 1.96 (the 97.5th percentile), to show a meaningful number of sources on the plot for this pronounced effect. Sources with representational change z-scores above these thresholds were cross-referenced to the Harvard-Oxford atlas [110] to assign labels based on their MNI coordinates. Sources where MNI coordinates did not have a label were allocated to the nearest label. As with our previous analyses, only the conditions whose distribution *p-*value (at the sensor level) demonstrated evidence for representational merging (i.e., *p*<.025) were modelled.

#### Behaviour-neural modelling at the session-level

To test the extent to which the changes in representational similarity across the intervention predicted the changes in subjects’ propensity to endorse similar lures, we conducted linear models at the session level (i.e., per visit). For this modelling, we calculated the degree of change in the subject’s propensity to endorse similar-object lures beyond different-object lures, per session across the intervention period. The score was calculated by taking a difference between the percentage accuracy for similar lures (correctly identifying a similar lure as new) and different lures (correctly identifying a different object association as new). This difference score was calculated for immediate recognition and for delayed recognition, per session. Using these similar-different accuracy scores, we computed the difference between the delayed recognition phase and the immediate recognition phase to quantify the change in subjects’ relative propensity to endorse similar lures erroneously as “old” (i.e., similar-different change score). The full calculation of similar-different change scores can be represented as: (delayed recognition similar-lure accuracy – delayed recognition different-lure accuracy) – (immediate recognition similar-lure accuracy – immediate recognition different-lure accuracy). Thus, positive values indicate an improvement in discrimination of similar-lures over different-lures across the intervention, whereas negative numbers indicate increased discrimination of different-lures over similar-lures across the intervention.

To initially determine the impact of the intervention conditions on the similar-different change score (i.e., the change in similar-over different-lure accuracy, from immediate to delayed recognition), we conducted a linear mixed-effects model using the subjects’ similar-different change scores across the intervention as the outcome, with intervention condition (retrieval training, restudy, sleep, or wake) as a fixed effect, and subject ID as a random effect. To identify where the significant differences were between each condition, treatment contrast coding was applied to every intervention condition in the model.

To link these changes in recognition accuracy to representational changes, subjects’ similar-different change scores were modelled as a function of the sensor-level representational change z-scores in a linear regression. For this modelling, the representational change z-scores were averaged across the time window and frequency band where the effect was most prominent in previous linear modelling across the intervention. Additionally, the intervention conditions were modelled separately to avoid multicollinearity between the condition intervention and representational change z-scores, and only the conditions whose distribution *p-*value demonstrated evidence for representational merging (i.e., *p*<.025) were modelled.

#### Behaviour-neural modelling at the item-level

We also aimed to predict accuracy at identifying individual similar lures at delayed recognition, using the item-level changes in representations across the intervention. Due to the nature of the paradigm, subjects were rarely (0−2 times per condition) shown both an object as a similar-lure (to get the behavioural accuracy), and the original object in a different pair (to measure the phase similarity), as each pairing was only tested once. Therefore, we modified our phase similarity comparisons for this analysis and compared each object in delayed recognition to its similar object and all different objects in immediate recognition. These item-level z-scores were computed in the same way as the original representational similarity z-scores, at the sensor-level: We calculated a distribution of different-object phase similarities per-subject and bootstrapped from these sets 10,000 times to create a distribution of means. Then the item-level similar-object phase similarities were standardised using the mean and standard deviation of the different-object to create a z-score for each object. Finally, we averaged these values based on the broad time window and frequency band where the representational merging was most prominent in previous session-level modelling. These item-level representational change z-scores represent the phase similarity of an object post-intervention, to its similar object pre-intervention, beyond that of different-object phase similarities.

We then used these item-level representational change z-scores to predict accuracy for similar lures during delayed recognition. This modelling was conducted using a linear mixed-effects logistic regression to predict binary accuracy at delayed recognition as the outcome. The model’s predictor was the item-level representational change z-score, with subject ID and object included as random effects. Also, conditions were treated in separate models, and only the conditions whose distribution *p-*value demonstrated evidence for representational merging (i.e., *p*<.025) were modelled.

## Acknowledgements

The authors would like to thank all of the subjects for their time.

## Supporting information

**S1 File. Sample characteristics.**

**S2 Table. Recognition accuracy model output.**

**S3 File. Object-specific cluster results.**

**S4 File. Encoding shift outputs.**

**S5 File. Intervention shift outputs.**

**S6 File. Session-level behaviour modelling.**

**S7 Text. Screening questions.**

