## Supplementary material for "Phase similarity between similar objects indicates representational merging across retrieval training but not sleep": S1 File

S1 File. Sample characteristics.

S1 Table A. Questionnaire scores.

| Questionnaire | Mean (SD) | Range |
| --- | --- | --- |
| Pittsburgh Sleep Quality Index | 4.13 (1.11) | 1–5 |
| Morningness-Eveningness Questionnaire | 50.70 (10.31) | 36–73 |
| Flinders Handedness Survey | 9.70 (0.79) | 7–10 |

*SD* = standard deviation.

IAF and custom frequency band descriptives.

IAF: *M*=9.84, *SD*=0.73, range=8.50-11.19.

S1 Table B. Custom frequency ranges.

| Frequency band | Lower limit mean (SD) | Upper limit mean (SD) |
| --- | --- | --- |
| Theta | 4.03 (0.51) | 6.20 (0.74) |
| Alpha | 8.13 (0.96) | 12.40 (1.48) |
| Sigma | 12.40 (1.48) | 16.25 (1.93) |
| Beta | 16.25 (1.93) | 24.80 (2.96) |

S1 Table C. Sleep intervention characteristics.

| Sleep Measure | Mean (SD) | Range |
| --- | --- | --- |
| TST (min) | 82.44 (26.31) | 25.00–116.00 |
| SOL (min) | 11.90 (11.11) | 1.50–53.00 |
| N1 (%) | 16.70 (9.54) | 5.00–49.50 |
| N2 (%) | 43.90 (14.00) | 19.50–71.00 |
| SWS (%) | 19.00 (18.31) | 0.00–56.00 |
| REM (%) | 2.87 (6.30) | 0.00–20.50 |

TST = total sleep time, SOL = sleep onset latency, N1 = stage 1 non-rapid eye movement (NREM) sleep, N2 = stage 2 NREM sleep, SWS = slow-wave sleep, REM = rapid eye movement sleep. No subject experienced sleep onset REM.
