## Supplementary material for "Phase similarity between similar objects indicates representational merging across retrieval training but not sleep": S2 Table

S2 Table. Recognition accuracy model output.

Recognition accuracy change score ~ condition * object_type + (1|ID)

| Effect | *χ^2^* | *df* | *p* |
| --- | --- | --- | --- |
| condition | 56.31 | 3 | <.001 |
| object_type | 5.27 | 2 | .072 |
| condition:object_type | 35.49 | 6 | <.001 |

*df* = degrees of freedom.
