## Supplementary material for "Phase similarity between similar objects indicates representational merging across retrieval training but not sleep": S3 File

**S3 File. Object-specific cluster results.**

For the first of our EEG analyses, we wanted to identify the times, frequencies, and channels contributing to object-specific representations from learning to immediate recognition. To do this, we first conducted a cluster-based permutation on the phase similarity for the same vs. different object comparison, to capture representations unique to objects. One significant cluster was detected where EEG phase was significantly more similar between same-objects compared to different-objects between learning and immediate recognition (*p*=.001). This cluster, visualised in the subfigure a, involved phase similarity in the frequencies 2−15 Hz, 20−580 ms post-object onset, across parietal and occipital channels. Next, we conducted another cluster-based permutation for the same-similar object comparison to detect a cluster of representations unique to object, beyond the conceptual likeness they share with their MST similar lures. This revealed one significant cluster (see subfigure c) where phase similarity was greater for same-object comparisons over similar-object comparisons between learning and immediate recognition (*p*=.036). This cluster encompassed frequencies spanning 2−27 Hz, across the entire epoch (−100−700 ms), with a broad distribution of channels across the entire scalp.

S3 Fig A. Same-different and same-similar object phase similarity clusters.

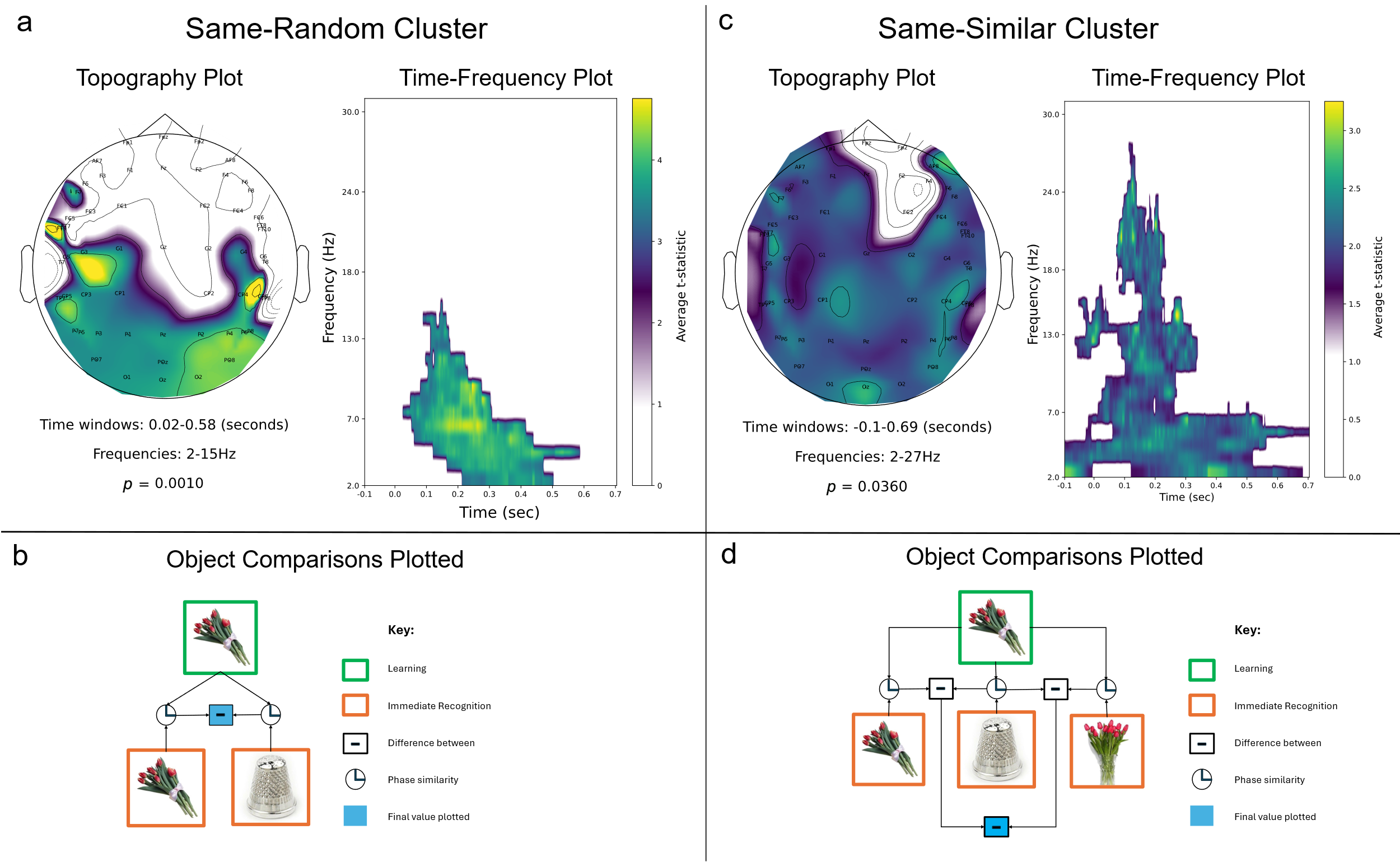

a) This section depicts the cluster of phase similarity scores between learning and immediate recognition comparison of same and different objects. The statistic was computed from a cluster-based permutation (forming clusters across time, frequencies, and channels). The topography plot shows the channels contributing to the cluster, averaged over times and frequencies. The time-frequency plot shows the times and frequencies contributing to the cluster, averaged over channels. b) Illustration of the same-different object comparison performed in the representational similarity analysis plotted in subfigure a. c) Cluster of phase similarity difference between same and similar objects. The statistic was computed from a cluster-based permutation (forming clusters across time, frequencies, and channels). The topography plot shows the channels contributing to the cluster, averaged over times and frequencies. The time-frequency plot shows the times and frequencies contributing to the cluster, averaged over channels. d) Illustration of the same-similar object comparison underlying subfigure c.
