## Supplementary material for "Phase similarity between similar objects indicates representational merging across retrieval training but not sleep": S4 File

**S4 File. Encoding shift outputs.**

S4 Table A. Linear model output – encoding shift.

phase_diff ~ band * time_win + (1|ID) + (1|ch_name)

| Effect | *χ^2^* | *df* | *p* |
| --- | --- | --- | --- |
| band | 92.77 | 3 | <.001 |
| time_win | 10.09 | 1 | .001 |
| band:time_win | 28.25 | 3 | <.001 |

*df* = degrees of freedom.

S4 Fig A. Same-different and same-similar object phase similarity clusters, for the average referenced data.

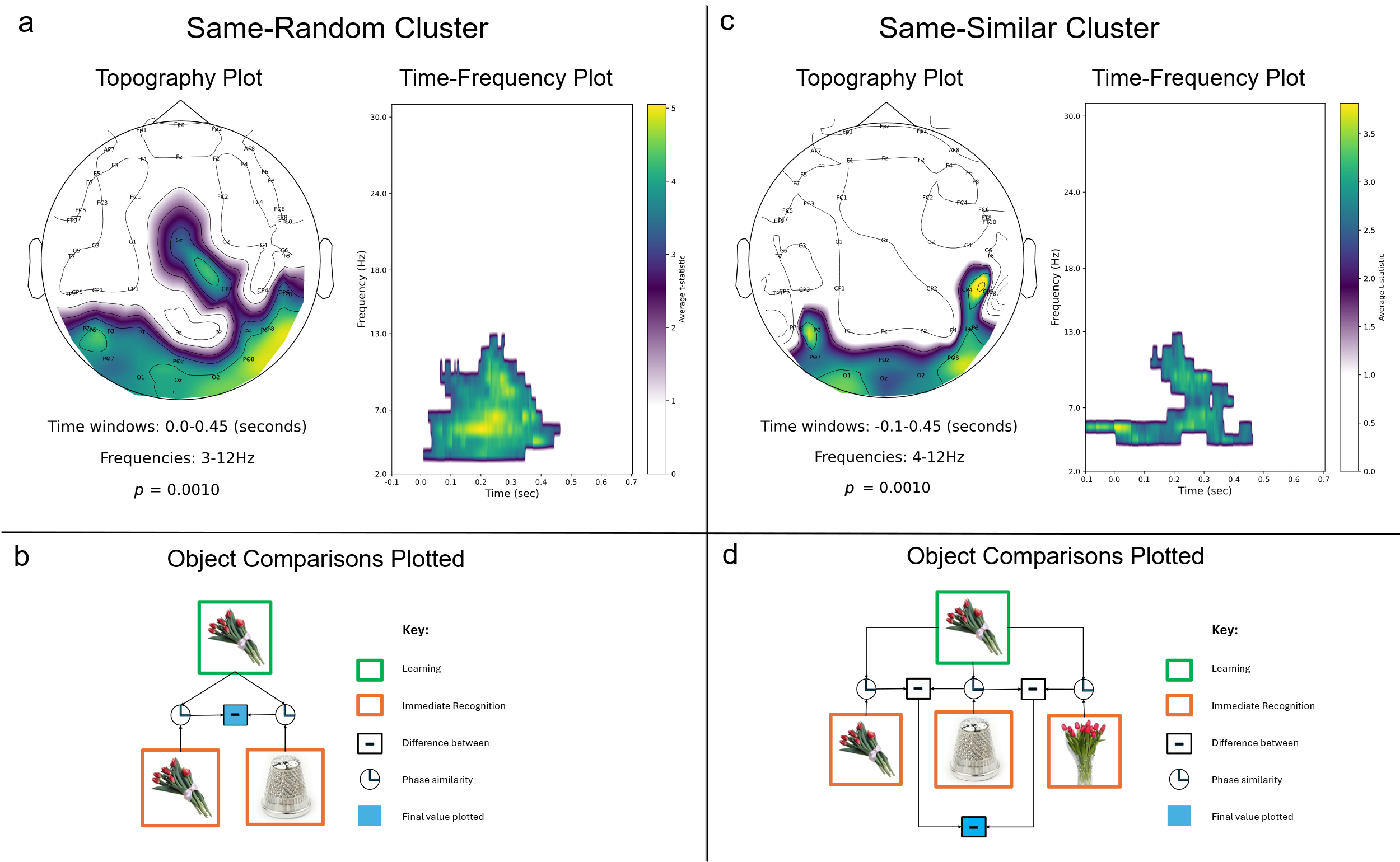

a) This section depicts the cluster of phase similarity difference scores determined from the cluster-based permutation across time, frequencies, and topography, for the comparison of same and different objects. The topography plot shows the channels contributing to the cluster, averaged over time and frequencies. The time-frequency plot shows the times and frequencies contributing to the cluster, averaged over channels. b) This key is a simplification of the calculations performed to get the z-scores that are plotted in the cluster graphs, for the same-different object comparison. c) This section depicts the cluster of phase similarity difference scores determined from the cluster-based permutation across time, frequencies, and topography, for the comparison of same and similar objects. The topography plot shows the channels contributing to the cluster, averaged over time and frequencies. The time-frequency plot shows the times and frequencies contributing to the cluster, averaged over channels. d) This key is a simplification of the object comparisons performed to get the z-scores that are plotted in the cluster graphs, for the same-similar object comparison.

S4 Fig B. Representational merging from learning to immediate recognition, for the average referenced data.

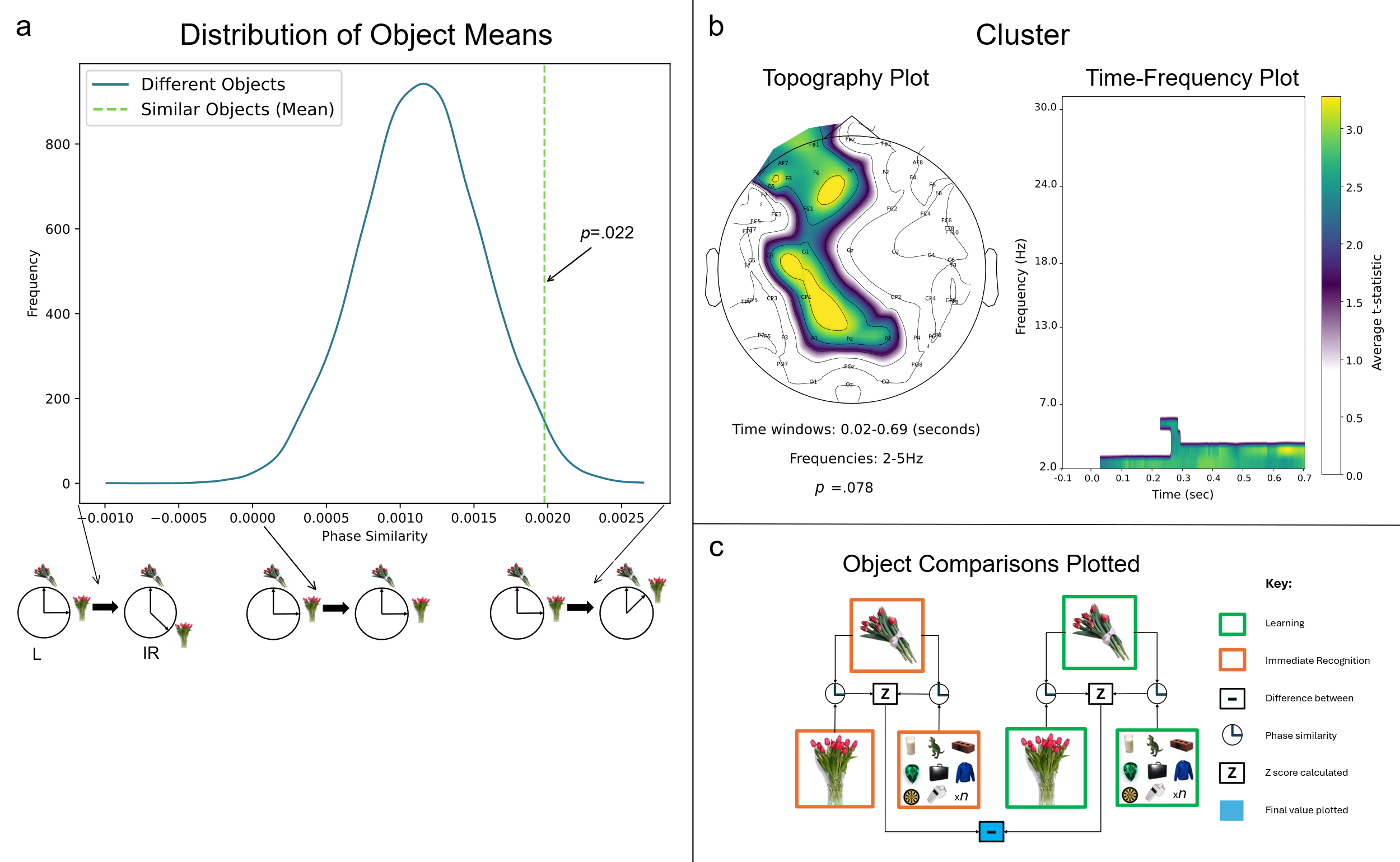

L = learning, IR = immediate recognition. a) The ratio highlighted in the distribution of object means refers to the ratio of different-object means that fall above the similar-object mean. The arrows and diagrams along the x-axis of the distribution graph graphically depict the direction of change in phase similarity that occurs from learning to immediate recognition. Negative values indicate decreased phase similarity of similar objects at immediate recognition, positive values indicate increased phase similarity of similar objects at immediate recognition, and values closer to 0 indicate little change from learning to immediate recognition. b) This section depicts the lowest *p*-value cluster of representational change z-scores determined from the cluster-based permutation across time, frequencies, and topography, for the encoding shift. The topography plot shows the channels contributing to the same cluster, averaged over time and frequencies. The time-frequency plot shows the times and frequencies contributing to the cluster, averaged over channels. c) This key is a simplification of the object comparisons performed to get the z-scores that are plotted in the cluster plots.

S4 Table B. The sources contributing to representational merging in the encoding shift.

| Region | Z-score |
| --- | --- |
| Cuneal Cortex* | 3.610286 |
| Occipital Pole | 3.433389 |
| Occipital Pole | 3.368327 |
| Cuneal Cortex | 3.287755 |
| Cuneal Cortex | 3.284609 |
| Precuneous Cortex* | 3.283758 |
| Cuneal Cortex | 3.23431 |
| Precuneous Cortex | 3.175432 |
| Lateral Occipital Cortex, superior division* | 3.088707 |
| Precuneous Cortex | 3.048152 |
| Occipital Pole | 3.015727 |
| Cuneal Cortex | 2.990822 |
| Lateral Occipital Cortex, superior division* | 2.952351 |
| Precuneous Cortex | 2.937046 |
| Lateral Occipital Cortex, superior division | 2.90953 |
| Lingual Gyrus | 2.902009 |
| Temporal Occipital Fusiform Cortex* | 2.883048 |
| Lateral Occipital Cortex, superior division | 2.868235 |
| Occipital Pole | 2.857403 |
| Lateral Occipital Cortex, superior division* | 2.844147 |
| Precuneous Cortex | 2.80497 |
| Intracalcarine Cortex | 2.793041 |
| Occipital Pole | 2.777427 |
| Cuneal Cortex | 2.759317 |
| Temporal Occipital Fusiform Cortex* | 2.755286 |
| Precuneous Cortex | 2.738626 |
| Lateral Occipital Cortex, superior division | 2.706178 |
| Temporal Occipital Fusiform Cortex* | 2.696939 |
| Occipital Fusiform Gyrus* | 2.681206 |
| Intracalcarine Cortex* | 2.674155 |
| Lateral Occipital Cortex, superior division | 2.652494 |
| Supracalcarine Cortex | 2.645354 |
| Angular Gyrus* | 2.63934 |
| Precentral Gyrus | 2.622412 |
| Lateral Occipital Cortex, superior division | 2.609404 |
| Occipital Pole* | 2.599115 |
| Occipital Fusiform Gyrus* | 2.595404 |
| Occipital Fusiform Gyrus* | 2.595057 |
| Lateral Occipital Cortex, superior division | 2.592317 |
| Occipital Pole* | 2.582396 |
| Lateral Occipital Cortex, superior division | 2.579731 |
| Occipital Fusiform Gyrus* | 2.579388 |
| Precuneous Cortex | 2.578065 |
| Lateral Occipital Cortex, inferior division* | 2.574267 |
| Cingulate Gyrus, posterior division* | 2.573091 |
| Cingulate Gyrus, posterior division* | 2.569154 |
| Superior Frontal Gyrus | 2.566944 |
| Precuneous Cortex | 2.559386 |
| Lateral Occipital Cortex, superior division | 2.55751 |
| Precentral Gyrus | 2.55312 |
| Precuneous Cortex | 2.551836 |
| Occipital Fusiform Gyrus* | 2.549871 |
| Precuneous Cortex | 2.545103 |
| Occipital Pole | 2.527269 |
| Precentral Gyrus | 2.523482 |
| Precuneous Cortex | 2.520608 |
| Occipital Fusiform Gyrus* | 2.518736 |
| Precuneous Cortex | 2.515015 |
| Intracalcarine Cortex | 2.48857 |
| Intracalcarine Cortex | 2.487951 |
| Precuneous Cortex | 2.480163 |
| Occipital Pole | 2.479822 |
| Temporal Occipital Fusiform Cortex* | 2.472744 |
| Lingual Gyrus | 2.47223 |
| Temporal Occipital Fusiform Cortex* | 2.464579 |
| Intracalcarine Cortex | 2.460055 |
| Superior Parietal Lobule* | 2.443011 |
| Precentral Gyrus | 2.439102 |
| Temporal Occipital Fusiform Cortex | 2.434801 |
| Occipital Fusiform Gyrus* | 2.428142 |
| Frontal Orbital Cortex | 2.427346 |
| Occipital Fusiform Gyrus* | 2.42669 |
| Precuneous Cortex* | 2.424383 |
| Parietal Opercular Cortex* | 2.415987 |
| Precuneous Cortex* | 2.40812 |
| Occipital Fusiform Gyrus | 2.406016 |
| Occipital Pole | 2.405976 |
| Postcentral Gyrus | 2.403178 |
| Cuneal Cortex* | 2.402296 |
| Precuneous Cortex* | 2.397974 |
| Precuneous Cortex* | 2.394977 |
| Parietal Opercular Cortex* | 2.389892 |
| Lingual Gyrus* | 2.387962 |
| Occipital Pole | 2.385467 |
| Precuneous Cortex* | 2.385415 |
| Intracalcarine Cortex* | 2.381131 |
| Precuneous Cortex* | 2.373169 |
| Occipital Fusiform Gyrus* | 2.37148 |
| Lingual Gyrus | 2.370085 |
| Lingual Gyrus* | 2.348929 |
| Frontal Pole | 2.348588 |
| Occipital Fusiform Gyrus | 2.343224 |
| Occipital Fusiform Gyrus* | 2.338669 |
| Lateral Occipital Cortex, superior division | 2.334747 |
| Precuneous Cortex | 2.327487 |
| Temporal Occipital Fusiform Cortex* | 2.324217 |
| Intracalcarine Cortex | 2.321942 |
| Temporal Occipital Fusiform Cortex* | 2.315797 |
| Precuneous Cortex | 2.309127 |
| Parietal Opercular Cortex* | 2.308489 |
| Intracalcarine Cortex | 2.306691 |
| Lateral Occipital Cortex, inferior division | 2.30664 |
| Temporal Occipital Fusiform Cortex | 2.302886 |
| Lateral Occipital Cortex, superior division* | 2.301408 |
| Temporal Occipital Fusiform Cortex* | 2.301285 |
| Temporal Occipital Fusiform Cortex* | 2.299544 |
| Precuneous Cortex* | 2.297307 |
| Postcentral Gyrus | 2.297101 |
| Precuneous Cortex* | 2.29261 |
| Occipital Pole* | 2.289288 |
| Occipital Fusiform Gyrus* | 2.288041 |
| Occipital Pole* | 2.283177 |
| Occipital Pole* | 2.282057 |
| Lingual Gyrus | 2.280789 |
| Precuneous Cortex | 2.280109 |
| Angular Gyrus* | 2.274413 |
| Occipital Pole* | 2.273088 |
| Lingual Gyrus | 2.272555 |
| Precentral Gyrus | 2.272482 |
| Lingual Gyrus* | 2.269038 |
| Lateral Occipital Cortex, superior division | 2.267463 |
| Lateral Occipital Cortex, inferior division | 2.264558 |
| Supramarginal Gyrus, posterior division | 2.264343 |
| Temporal Occipital Fusiform Cortex* | 2.264127 |
| Superior Frontal Gyrus | 2.260452 |
| Precuneous Cortex | 2.259709 |
| Lingual Gyrus* | 2.257329 |
| Precuneous Cortex | 2.256884 |
| Occipital Pole | 2.254788 |
| Temporal Occipital Fusiform Cortex | 2.253465 |
| Occipital Pole | 2.249175 |
| Temporal Occipital Fusiform Cortex* | 2.248874 |
| Cingulate Gyrus, posterior division* | 2.245007 |
| Cingulate Gyrus, posterior division* | 2.244764 |
| Temporal Occipital Fusiform Cortex* | 2.237692 |
| Lingual Gyrus* | 2.232709 |
| Temporal Pole | 2.230007 |
| Occipital Pole | 2.22491 |
| Postcentral Gyrus | 2.21814 |
| Lingual Gyrus* | 2.217523 |
| Precentral Gyrus | 2.217469 |
| Intracalcarine Cortex* | 2.210054 |
| Superior Frontal Gyrus | 2.208882 |
| Superior Parietal Lobule | 2.207019 |
| Occipital Pole | 2.206722 |
| Parietal Opercular Cortex* | 2.206428 |
| Lingual Gyrus* | 2.197507 |
| Postcentral Gyrus | 2.195352 |
| Frontal Orbital Cortex* | 2.194345 |
| Central Opercular Cortex* | 2.193932 |
| Intracalcarine Cortex* | 2.19063 |
| Precuneous Cortex* | 2.187564 |
| Occipital Fusiform Gyrus* | 2.182645 |
| Occipital Fusiform Gyrus* | 2.181466 |
| Lateral Occipital Cortex, superior division* | 2.181196 |
| Superior Parietal Lobule | 2.180386 |
| Superior Temporal Gyrus, anterior division | 2.176113 |
| Precentral Gyrus | 2.175257 |
| Temporal Occipital Fusiform Cortex | 2.173441 |
| Intracalcarine Cortex | 2.17328 |
| Occipital Fusiform Gyrus | 2.170551 |
| Occipital Pole* | 2.169599 |
| Temporal Occipital Fusiform Cortex* | 2.166143 |
| Occipital Pole | 2.162568 |
| Superior Temporal Gyrus, anterior division | 2.160326 |
| Cuneal Cortex* | 2.159313 |
| Occipital Pole | 2.152942 |
| Angular Gyrus | 2.150757 |
| Lingual Gyrus | 2.148802 |
| Lingual Gyrus* | 2.148143 |
| Precentral Gyrus | 2.147429 |
| Lingual Gyrus* | 2.147265 |
| Angular Gyrus* | 2.146048 |
| Lateral Occipital Cortex, inferior division | 2.142894 |
| Angular Gyrus | 2.138664 |
| Temporal Occipital Fusiform Cortex* | 2.135801 |
| Postcentral Gyrus | 2.133471 |
| Frontal Orbital Cortex | 2.130683 |
| Parietal Opercular Cortex | 2.129339 |
| Intracalcarine Cortex | 2.120218 |
| Lateral Occipital Cortex, superior division | 2.119805 |
| Cuneal Cortex | 2.11571 |
| Precuneous Cortex* | 2.108865 |
| Superior Parietal Lobule* | 2.106325 |
| Lingual Gyrus* | 2.104742 |
| Occipital Pole | 2.103745 |
| Precentral Gyrus | 2.097006 |
| Lateral Occipital Cortex, inferior division* | 2.096444 |
| Precuneous Cortex | 2.095755 |
| Postcentral Gyrus | 2.093352 |
| Lingual Gyrus* | 2.092496 |
| Lingual Gyrus | 2.091939 |
| Lingual Gyrus* | 2.090645 |
| Occipital Fusiform Gyrus* | 2.089395 |
| Angular Gyrus* | 2.089243 |
| Lateral Occipital Cortex, inferior division | 2.089189 |
| Planum Temporale* | 2.08665 |
| Superior Parietal Lobule | 2.080675 |
| Lingual Gyrus | 2.079484 |
| Temporal Occipital Fusiform Cortex* | 2.077324 |
| Precentral Gyrus | 2.072901 |
| Lingual Gyrus* | 2.072339 |
| Lingual Gyrus* | 2.066172 |
| Lateral Occipital Cortex, inferior division | 2.065183 |
| Temporal Occipital Fusiform Cortex | 2.064712 |
| Lateral Occipital Cortex, inferior division | 2.063258 |
| Central Opercular Cortex | 2.062391 |
| Lingual Gyrus | 2.060863 |
| Superior Frontal Gyrus | 2.060813 |
| Lingual Gyrus | 2.059546 |
| Occipital Pole* | 2.058876 |
| Precuneous Cortex | 2.057173 |
| Temporal Occipital Fusiform Cortex* | 2.056761 |
| Lingual Gyrus | 2.055115 |
| Occipital Fusiform Gyrus | 2.054865 |
| Temporal Pole | 2.050422 |
| Parietal Opercular Cortex | 2.049772 |
| Lingual Gyrus* | 2.047438 |
| Lateral Occipital Cortex, superior division* | 2.044793 |
| Lateral Occipital Cortex, superior division | 2.044502 |
| Temporal Occipital Fusiform Cortex | 2.043955 |
| Planum Polare | 2.038782 |
| Occipital Pole | 2.032952 |
| Temporal Occipital Fusiform Cortex* | 2.032297 |
| Insular Cortex | 2.030444 |
| Occipital Fusiform Gyrus | 2.02996 |
| Occipital Fusiform Gyrus | 2.028746 |
| Planum Temporale* | 2.027966 |
| Occipital Pole* | 2.026629 |
| Occipital Fusiform Gyrus* | 2.026454 |
| Lateral Occipital Cortex, inferior division | 2.026186 |
| Temporal Occipital Fusiform Cortex* | 2.025631 |
| Superior Parietal Lobule* | 2.024665 |
| Occipital Fusiform Gyrus* | 2.023766 |
| Lateral Occipital Cortex, superior division* | 2.021735 |
| Lateral Occipital Cortex, superior division | 2.020949 |
| Occipital Pole* | 2.019448 |
| Cuneal Cortex | 2.019361 |
| Planum Polare | 2.018153 |
| Supramarginal Gyrus, anterior division | 2.016531 |
| Temporal Occipital Fusiform Cortex* | 2.016127 |
| Cingulate Gyrus, posterior division | 2.015509 |
| Lateral Occipital Cortex, inferior division | 2.015105 |
| Lateral Occipital Cortex, inferior division* | 2.013263 |
| Temporal Pole | 2.012554 |
| Occipital Pole* | 2.010403 |
| Precuneous Cortex* | 2.007425 |
| Lateral Occipital Cortex, superior division | 2.006966 |
| Intracalcarine Cortex | 2.006259 |
| Angular Gyrus | 2.004725 |
| Precentral Gyrus | 2.003792 |
| Lateral Occipital Cortex, inferior division | 2.003447 |
| Lingual Gyrus* | 2.001599 |
| Precuneous Cortex | 2.000541 |
| Temporal Occipital Fusiform Cortex* | 1.99621 |
| Postcentral Gyrus | 1.995835 |
| Lateral Occipital Cortex, inferior division* | 1.994476 |
| Occipital Fusiform Gyrus* | 1.990068 |
| Cingulate Gyrus, posterior division* | 1.988349 |
| Superior Temporal Gyrus, anterior division | 1.985597 |
| Lateral Occipital Cortex, inferior division* | 1.984651 |
| Temporal Occipital Fusiform Cortex* | 1.9838 |
| Occipital Fusiform Gyrus* | 1.983216 |
| Occipital Pole* | 1.981961 |
| Occipital Fusiform Gyrus | 1.979451 |
| Temporal Occipital Fusiform Cortex* | 1.977084 |
| Precentral Gyrus | 1.974328 |
| Lateral Occipital Cortex, superior division | 1.973814 |
| Occipital Fusiform Gyrus* | 1.973303 |
| Superior Parietal Lobule | 1.971548 |
| Lateral Occipital Cortex, superior division | 1.970603 |
| Postcentral Gyrus* | 1.968411 |
| Lateral Occipital Cortex, superior division | 1.968166 |
| Lateral Occipital Cortex, inferior division | 1.963858 |
| Lingual Gyrus* | 1.963407 |

* = sources that were assigned the nearest non-background label. The labels were taken from the Harvard-Oxford dictionary, based on the MNI coordinates of the source. The representational change z-scores of each source are all above the 97.5th percentile and are averaged over times and frequencies in the encoding cluster.
