## Supplementary material for "Phase similarity between similar objects indicates representational merging across retrieval training but not sleep": S5 File

**S5 File. Intervention shift outputs.**

S5 Table A. Linear model output – intervention shift.

phase_diff ~ band * time_win + (1|ID) + (1|ch_name)

| Effect | *F* | *df* | *p* |
| --- | --- | --- | --- |
| band | 116.38 | 3 | <.001 |
| time_win | 4.06 | 1 | .044 |
| band:time_win | 34.06 | 3 | <.001 |

*df* = degrees of freedom.

S5 Fig A. Representational merging from immediate to delayed recognition, for the average referenced data**.**

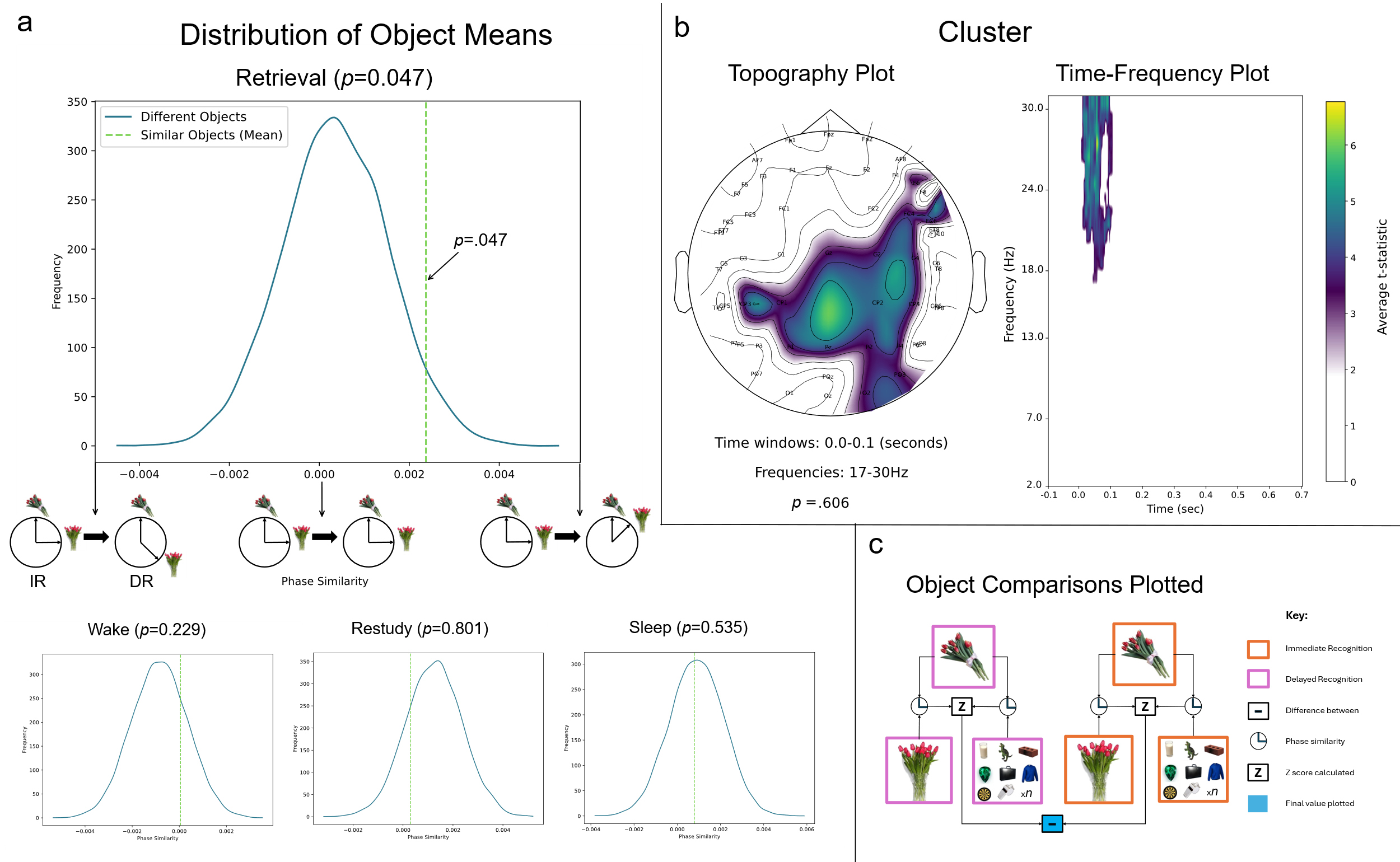

IR = immediate recognition, DR = delayed recognition. a) The ratio highlighted in the distribution of object means refers to the ratio of different-object means that fall above the similar-object mean. The arrows and diagrams along the x-axis of the distribution graph graphically depict the direction of change in phase similarity that occurs from immediate recognition to delayed recognition. Negative values indicate decreased phase similarity of similar objects at delayed recognition, positive values indicate increased phase similarity of similar objects at delayed recognition, and values closer to 0 indicate little change from immediate to delayed recognition. b) This section depicts the lowest *p*-values cluster of representational change z-scores determined from the cluster-based permutation across time, frequencies, and topography, for the intervention shift. The topography plot shows the channels contributing to this cluster, averaged over time and frequencies. The time-frequency plot shows the times and frequencies contributing to the cluster, averaged over channels. c) This key is a simplification of the object comparisons performed to get the z-scores that are plotted in the cluster plots.

S5 Table B. The sources contributing to representational merging in the retrieval training’s intervention shift.

| Region | Z-score |
| --- | --- |
| Lateral Occipital Cortex, inferior division | 2.445826 |
| Parietal Opercular Cortex* | 2.427392 |
| Parahippocampal Gyrus, posterior division* | 2.331251 |
| Lingual Gyrus* | 2.288233 |
| Supramarginal Gyrus, anterior division | 2.287396 |
| Inferior Temporal Gyrus, posterior division | 2.285315 |
| Parietal Opercular Cortex* | 2.275704 |
| Inferior Temporal Gyrus, temporooccipital part* | 2.273214 |
| Inferior Temporal Gyrus, posterior division | 2.267414 |
| Parietal Opercular Cortex | 2.256483 |
| Lingual Gyrus | 2.213528 |
| Parahippocampal Gyrus, posterior division* | 2.211534 |
| Lateral Occipital Cortex, inferior division | 2.187111 |
| Lingual Gyrus* | 2.158867 |
| Parietal Opercular Cortex | 2.156303 |
| Lingual Gyrus | 2.123423 |
| Planum Temporale | 2.081967 |
| Temporal Fusiform Cortex, posterior division | 2.046327 |
| Intracalcarine Cortex | 2.045143 |
| Temporal Fusiform Cortex, posterior division | 2.036757 |
| Inferior Temporal Gyrus, temporooccipital part* | 2.031056 |
| Planum Temporale* | 2.013499 |
| Frontal Pole | 2.009094 |
| Postcentral Gyrus | 2.005623 |
| Superior Temporal Gyrus, posterior division | 2.005462 |
| Superior Parietal Lobule* | 1.994609 |
| Lingual Gyrus | 1.992909 |
| Inferior Temporal Gyrus, temporooccipital part* | 1.987299 |
| Occipital Fusiform Gyrus* | 1.981476 |
| Postcentral Gyrus | 1.981282 |
| Postcentral Gyrus* | 1.980683 |
| Occipital Fusiform Gyrus | 1.978575 |
| Parietal Opercular Cortex* | 1.977752 |
| Lingual Gyrus* | 1.975243 |
| Supramarginal Gyrus, anterior division* | 1.972651 |
| Inferior Temporal Gyrus, temporooccipital part* | 1.971383 |
| Angular Gyrus* | 1.961389 |
| Supramarginal Gyrus, anterior division* | 1.957754 |
| Parahippocampal Gyrus, posterior division* | 1.946544 |
| Lingual Gyrus | 1.944044 |
| Lingual Gyrus* | 1.930077 |
| Occipital Fusiform Gyrus* | 1.929982 |
| Heschl's Gyrus (includes H1 and H2)* | 1.929654 |
| Frontal Pole | 1.929601 |
| Occipital Pole | 1.928322 |
| Superior Parietal Lobule | 1.924446 |
| Supramarginal Gyrus, posterior division | 1.918625 |
| Parahippocampal Gyrus, posterior division* | 1.918277 |
| Lateral Occipital Cortex, inferior division | 1.917294 |
| Central Opercular Cortex* | 1.916706 |
| Temporal Fusiform Cortex, posterior division* | 1.909189 |
| Inferior Temporal Gyrus, posterior division* | 1.902406 |
| Parietal Opercular Cortex | 1.888452 |
| Temporal Occipital Fusiform Cortex* | 1.880122 |
| Intracalcarine Cortex | 1.877455 |
| Occipital Fusiform Gyrus* | 1.875197 |
| Lingual Gyrus* | 1.871478 |
| Parahippocampal Gyrus, posterior division* | 1.870345 |
| Temporal Occipital Fusiform Cortex* | 1.867681 |
| Lingual Gyrus* | 1.865232 |
| Occipital Pole* | 1.859396 |
| Frontal Pole | 1.852865 |
| Lingual Gyrus | 1.851399 |
| Lingual Gyrus* | 1.839977 |
| Planum Polare* | 1.838727 |
| Lingual Gyrus | 1.834542 |
| Temporal Occipital Fusiform Cortex | 1.833787 |
| Occipital Fusiform Gyrus* | 1.833051 |
| Precentral Gyrus | 1.829572 |
| Inferior Temporal Gyrus, posterior division* | 1.828496 |
| Lingual Gyrus* | 1.826681 |
| Inferior Temporal Gyrus, posterior division | 1.820967 |
| Occipital Fusiform Gyrus* | 1.81754 |
| Lingual Gyrus | 1.817463 |
| Occipital Fusiform Gyrus* | 1.810418 |
| Intracalcarine Cortex | 1.808598 |
| Occipital Fusiform Gyrus | 1.798365 |
| Occipital Fusiform Gyrus | 1.797566 |
| Lateral Occipital Cortex, superior division* | 1.79659 |
| Occipital Fusiform Gyrus* | 1.79543 |
| Parahippocampal Gyrus, posterior division | 1.792443 |
| Inferior Temporal Gyrus, temporooccipital part | 1.789034 |
| Occipital Fusiform Gyrus* | 1.788195 |
| Occipital Pole* | 1.780354 |
| Parahippocampal Gyrus, posterior division* | 1.779156 |
| Occipital Fusiform Gyrus* | 1.776932 |
| Lateral Occipital Cortex, inferior division* | 1.772577 |
| Parahippocampal Gyrus, posterior division | 1.770781 |
| Lateral Occipital Cortex, inferior division | 1.770607 |
| Occipital Pole | 1.767734 |
| Lingual Gyrus | 1.766744 |
| Occipital Fusiform Gyrus* | 1.765145 |
| Occipital Fusiform Gyrus* | 1.762489 |
| Lateral Occipital Cortex, inferior division* | 1.761022 |
| Heschl's Gyrus (includes H1 and H2) | 1.760993 |
| Temporal Occipital Fusiform Cortex | 1.760453 |
| Occipital Fusiform Gyrus* | 1.756024 |
| Temporal Fusiform Cortex, posterior division* | 1.752779 |
| Occipital Pole | 1.750047 |
| Temporal Occipital Fusiform Cortex* | 1.744751 |
| Planum Temporale* | 1.741924 |
| Lateral Occipital Cortex, inferior division | 1.740078 |
| Supramarginal Gyrus, anterior division* | 1.73724 |
| Inferior Temporal Gyrus, temporooccipital part* | 1.731544 |
| Occipital Fusiform Gyrus* | 1.727756 |
| Lingual Gyrus | 1.725413 |
| Postcentral Gyrus | 1.722821 |
| Lingual Gyrus | 1.722529 |
| Superior Parietal Lobule | 1.721437 |
| Occipital Fusiform Gyrus* | 1.718859 |
| Occipital Pole* | 1.713911 |
| Lingual Gyrus* | 1.70741 |
| Middle Temporal Gyrus, temporooccipital part | 1.702911 |
| Temporal Occipital Fusiform Cortex* | 1.7015 |
| Temporal Occipital Fusiform Cortex* | 1.698469 |
| Lingual Gyrus* | 1.695383 |
| Occipital Pole* | 1.695002 |
| Occipital Pole | 1.692981 |
| Occipital Pole* | 1.691381 |
| Planum Temporale | 1.688459 |
| Supramarginal Gyrus, anterior division | 1.687475 |
| Middle Temporal Gyrus, temporooccipital part | 1.687317 |
| Lateral Occipital Cortex, inferior division* | 1.686277 |
| Lingual Gyrus* | 1.685822 |
| Occipital Fusiform Gyrus | 1.6844 |
| Lingual Gyrus* | 1.682082 |
| Parahippocampal Gyrus, posterior division* | 1.680892 |
| Inferior Temporal Gyrus, temporooccipital part* | 1.6796 |
| Lingual Gyrus* | 1.679082 |
| Temporal Occipital Fusiform Cortex* | 1.676162 |
| Intracalcarine Cortex* | 1.675666 |
| Occipital Pole | 1.672823 |
| Supramarginal Gyrus, anterior division* | 1.672066 |
| Lateral Occipital Cortex, inferior division | 1.671601 |
| Lingual Gyrus* | 1.669688 |
| Lingual Gyrus* | 1.667106 |
| Temporal Occipital Fusiform Cortex* | 1.666532 |
| Temporal Fusiform Cortex, posterior division* | 1.664836 |
| Lingual Gyrus* | 1.662175 |
| Parahippocampal Gyrus, anterior division* | 1.661382 |
| Parahippocampal Gyrus, posterior division* | 1.66135 |
| Occipital Pole | 1.661072 |
| Lingual Gyrus* | 1.659795 |
| Parahippocampal Gyrus, posterior division* | 1.65873 |
| Occipital Pole* | 1.656698 |
| Occipital Pole | 1.656061 |
| Postcentral Gyrus | 1.652582 |
| Intracalcarine Cortex* | 1.651714 |
| Temporal Occipital Fusiform Cortex* | 1.650413 |
| Occipital Fusiform Gyrus* | 1.648898 |
| Cingulate Gyrus, posterior division* | 1.647916 |
| Occipital Fusiform Gyrus* | 1.646271 |

* = sources that were assigned the nearest non-background label. The labels were taken from the Harvard-Oxford dictionary, based on the MNI coordinates of the source. The representational change z-scores of each source are all above the 95th percentile and are averaged 400–700 ms and 7–13 Hz.
