## Supplementary material for "Phase similarity between similar objects indicates representational merging across retrieval training but not sleep": S6 File

S6 File. Session-level behaviour modelling.

We aimed to test if the change in propensity for subjects to endorse similar lures from pre- to post-intervention, differed based on the intervention condition. To address this, we conducted a linear mixed-effects model predicting subjects’ similar-different change scores (i.e., the change in similar- over different-lure accuracy, from immediate to delayed recognition), using the intervention condition as a fixed effect. This model revealed a significant effect of condition, *χ^2^*(3)=18.47, *p<*.001, which is displayed in Fig A. This figure illustrates that the propensity to endorse similar lures more than different lures generally increased across the retrieval training and wake interventions, but decreased across the sleep and restudy periods. Treatment contrast coding revealed that retrieval training and sleep conditions did not differ significantly.

S6 Fig A. Condition differences for the change in similar-lure over different-lure accuracy.

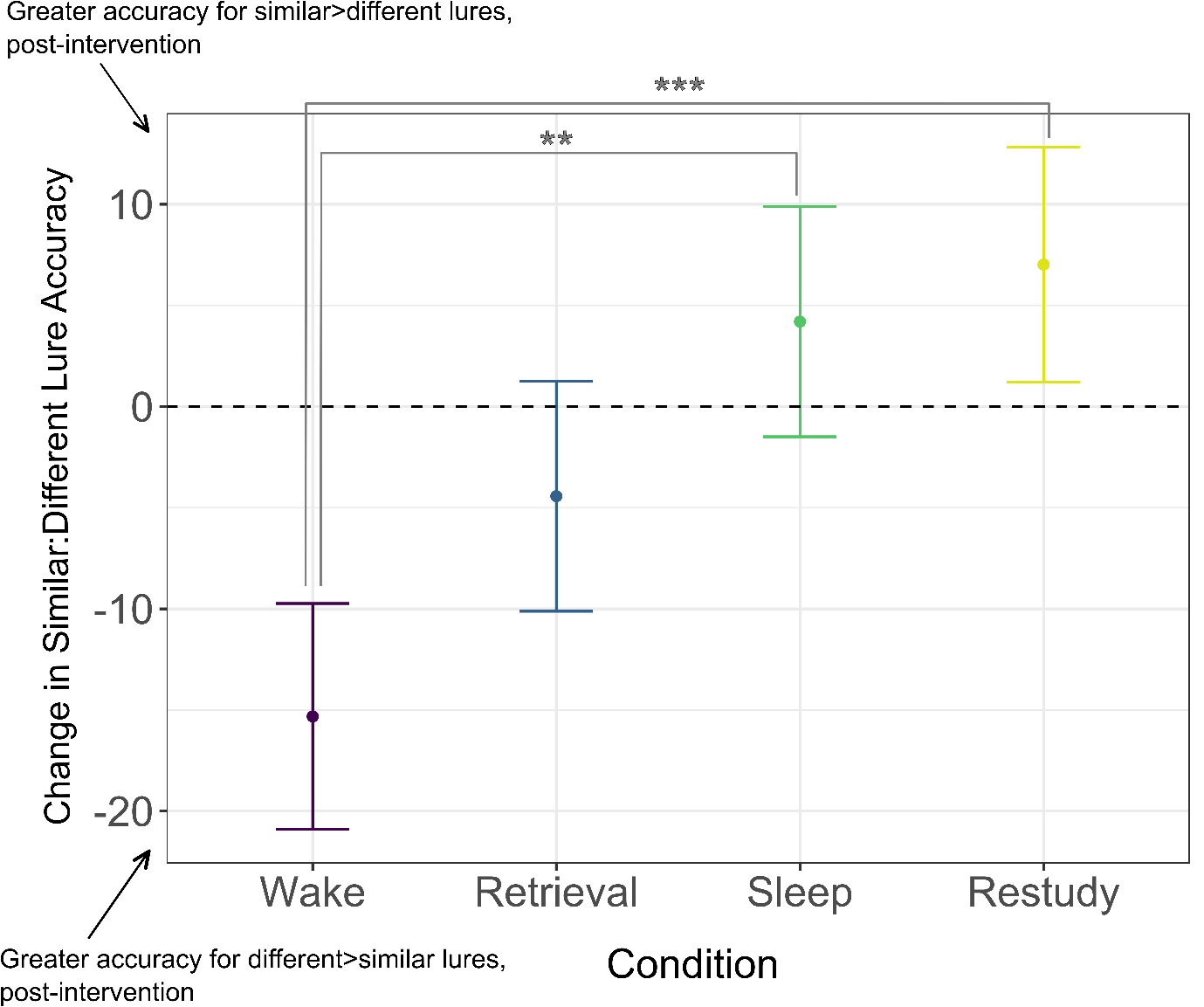
 The y-axis represents the difference between similar-lure accuracy and different-lure accuracy, from immediate to delayed recognition. Positive numbers indicate that there was a greater accuracy for similar- over different-object lures post-intervention, compared to pre-intervention. Negative numbers indicate that there was a greater accuracy for different over similar lures post-intervention, compared to pre-intervention. The zero point is marked with a dotted line, indicating little to no change in the difference between similar- over different lure accuracy across the intervention. The error bars represent the 83% confidence intervals. Significant contrasts are marked as: *denotes *p*<.05, ***p*<.01, and ****p*<.001.

Additionally, we used the representational change z-scores (averaged over the 400–700ms alpha-band) to predict subjects’ similar-different change scores (i.e., the change in session-level endorsement of similar lures over different lures, across the intervention period), for the retrieval training intervention. The purpose of this was to have an analogous behaviour measure of representational change to our neural measure. The linear model regression predicting similar-different change scores across retrieval training from the representational change z-scores, did not yield statistical significance *F*(1)=0.004, *p*=.950.
