## Supplementary material for "Phase similarity between similar objects indicates representational merging across retrieval training but not sleep": S7 Text

S7 Text. Screening questions.

1. What is your gender? (Woman/Man/Non-Binary/None of the above)
2. What is your age in years?
3. Are you a fluent English speaker? (Yes/No)
4. Do you have a history of, or any diagnosed, psychiatric or sleep illnesses (e.g., depression, ADHD, insomnia)? (Yes/No)
5. Have you taken recreational drugs in the last 6 months? (Yes/No)
6. Do you currently take any medication? If so, please enter the name/s. (Yes/No)
7. Do you currently take any medication? If so, please enter the name/s. (Yes/No)
8. Do you have normal/corrected vision? (Yes/No)
9. Do you have normal/corrected hearing? (Yes/No)
